# Culturable diversity of Arctic phytoplankton during pack ice melting

**DOI:** 10.1101/642264

**Authors:** Catherine Gérikas Ribeiro, Adriana Lopes dos Santos, Priscillia Gourvil, Florence Le Gall, Dominique Marie, Margot Tragin, Ian Probert, Daniel Vaulot

**Affiliations:** Sorbonne Université, CNRS, UMR7144, Team ECOMAP, Station Biologique de Roscoff, Roscoff, France; GEMA Center for Genomics, Ecology & Environment, Universidad Mayor, Camino La Pirámide, 5750, Huechuraba, Santiago, Chile; Nanyang Technological University, Asian School of the Environment, Singapore; Sorbonne Université, CNRS, FR2424, Roscoff Culture Collection, Station Biologique de Roscoff, Roscoff, France

## Abstract

Massive phytoplankton blooms develop at the Arctic ice edge, sometimes extending far under the pack ice. An extensive culturing effort was conducted before and during a phytoplankton bloom in Baffin Bay between April and July 2016. Different isolation strategies were applied, including flow cytometry cell sorting, manual single cell pipetting and serial dilution. Although all three techniques yielded the most common organisms, each technique retrieved specific taxa, highlighting the importance of using several methods to maximize the number and diversity of isolated strains. More than 1,000 cultures were obtained, characterized by 18S rRNA sequencing and optical microscopy and de-replicated to a subset of 276 strains presented in this work. Strains grouped into 57 genotypes defined by 100% 18S rRNA sequence similarity. These genotypes spread across five divisions: Heterokontophyta, Chlorophyta, Cryptophyta, Haptophyta and Dinophyta. Diatoms were the most abundant group (193 strains), mostly represented by the genera *Chaetoceros* and *Attheya*. The genera *Rhodomonas* and *Pyramimonas* were the most abundant non-diatom nanoplankton strains, while *Micromonas polaris* dominated the picoplankton. Diversity at the class level was higher during the peak of the bloom. Potentially new species were isolated, in particular within the genera *Navicula*, *Nitzschia*, *Coscinodiscus*, *Thalassiosira*, *Pyramimonas*, *Mantoniella* and *Isochrysis*.

Submitted to: Elementa: Science of the Anthropocene Date: May 17, 2019

## Introduction

Polar algal communities impact (Lutz et al., 2016) and are impacted by (Brown and Arrigo, 2013) ice melting cycles. The tight links between phytoplankton diversity/production and sea ice dynamics are beginning to be decoded and seem to be fairly complex (Arrigo et al., 2014; Olsen et al., 2017). Due to increased light availability and vertical mixing, shrinking of pack ice and the shift from thick multi-year ice to thinner first-year ice has been linked to enhanced Arctic primary production (Arrigo et al., 2008; Brown and Arrigo, 2013). However, the salinity decrease in surface waters resulting from higher ice melting rates and increased river run off leads to an increase in water column stratification which in turn may impact nutrient availability and plankton diversity (Li et al., 2009). The presence of ice-associated algae may impact the quantity (Kohlbach et al., 2016; Leu et al., 2011) and quality (Duerksen et al., 2014; Schmidt et al., 2018) of secondary production at high latitudes, as well as the recruitment of ice-associated diatoms to the water column (Kauko et al., 2018). Climate-related changes can also increase Arctic vulnerability to invasive species (Vincent, 2010) as the intrusion of warmer waters “atlantifies” the Arctic Ocean (Årthun et al., 2012) and temperate phy-toplankton move northwards, replacing Arctic communities (Neukermans et al., 2018).

Massive Arctic phytoplankton blooms have recently been detected not only along the ice edge (Perrette et al., 2011), but also extending large distances under the pack ice (Arrigo et al., 2012). The increasingly thin pack ice and the formation of melt-ponds allow viable areas for sub-ice bloom formation in almost one third of the ice-covered Arctic Ocean in the summer (Horvat et al., 2017).

The Arctic phytoplankton community exhibits strong seasonal variability (Sherr et al., 2003), with smaller organisms (picoplankton) dominating during winter and early summer, followed by diatoms during the spring bloom (Marquardt et al., 2016). The green alga *Micromonas* (Mamiellophyceae) is recognized as the dominant picophytoplankton (0.2 – 3 *µ*m) genus during the Arctic summer (Lovejoy et al., 2007; Balzano et al., 2017). The genus *Micromonas* is widespread throughout the world oceans (Tragin and Vaulot, 2019; Worden et al., 2009) and genetically diversified (Simon et al., 2017) in relation to thermal niches (Demory et al., 2018), with one species, *Micromonas polaris*, restricted to polar regions. Another Mamiellophyceae, *Bathycoccus prasinos*, may replace *M. polaris* during polar winter (Joli et al., 2017).

Regarding the more diverse Arctic nano-phytoplankton, the genus *Pyramimonas* has often been reported (Joli et al., 2017; Lovejoy et al., 2002) displaying high intra-generic diversity (Balzano et al., 2012; Daugbjerg and Moestrup, 1993), often associated to the sea ice (Harðardóttir et al., 2014; Kauko et al., 2018). The mamiellophyte *Mantoniella* is reported to a lesser extent in Arctic waters (Joli et al., 2017; Lovejoy et al., 2007; Terrado et al., 2013), although diversity within this genus seems to be higher than other polar Mamiellophyceae (Yau et al., 2018). Other commonly observed Arctic taxa include the bloom-forming *Phaeocystis* (Assmy et al., 2017), unidentified Pelagophyceae, the mixotroph and cosmopolitan *Dinobryon faculiferum* and *Chlamydomonas* (Balzano et al., 2012; Crawford et al., 2018; Lovejoy et al., 2002; Terrado et al., 2013).

Large size classes of polar phytoplankton are dominated by diatoms and dinoflagellates (Crawford et al., 2018). Diatoms constitute a large fraction of polar phytoplankton communities, especially in coastal areas and during episodic upwelling (Arrigo et al., 2014; Sherr et al., 2003), impacting carbon flux to the benthic community (Booth et al., 2002). This group is particularly important during late spring/summer months in the pelagic environment (Balzano et al., 2012; Lovejoy et al., 2002), but also comprises a significant portion of protist biomass during young ice formation (Kauko et al., 2018). The genera most often reported among Arctic centric diatoms are *Chaetoceros* and *Thalassiosira* (Johnsen et al., 2018; Lovejoy et al., 2002). *Chaetoceros gelidus* and *Chaetoceros neogracilis* often dominate Arctic phytoplankton assemblages (Crawford et al., 2018; Katsuki et al., 2009), although other species such as *C. decipiens* or *C. furcellatus* have also been reported (Johnsen et al., 2018; Joo et al., 2012). *Thalassiosira* is a diverse genus with both Arctic, ice-associated and subpolar/temperate water representatives (Luddington et al., 2016), in particular *T. nordenskioeldii* and *T. antarctica* var. *borealis* (Johnsen et al., 2018; Lovejoy et al., 2002; Poulin et al., 2011). High abundances of pennate diatoms are linked to late autumn/winter sea ice (Niemi et al., 2011) and bottom communities (Kauko et al., 2018; Leeuwe et al., 2018). The most commonly reported genera include *Cylindrotheca*, *Fragilariopsis*, *Navicula*, *Nitzschia* and *Pseudo-nitzschia* (Katsuki et al., 2009; Leeuwe et al., 2018; Poulin et al., 2011). In contrast to diatoms and small flagellates that present a strong seasonal signal, dinoflagellates are prevalent throughout the year (Comeau et al., 2011; Marquardt et al., 2016), although some taxa vary seasonally (Onda et al., 2017).

Few extensive, large scale culturing efforts have been carried out in the Arctic, with the exception of the MALINA cruise in summer 2009, which covered the Northeast Pacific Ocean, the Bering Strait, the Chukchi and Beaufort Seas (Balzano et al., 2012; 2017). As microbial communities respond to the rapid loss in Arctic ice cover and thickness (Comeau et al., 2011; Vincent, 2010), it is important to continue to attempt to culture phytoplankton from the region in order to dispose of reference strains whose physiology and taxonomy can be studied in the laboratory under controlled conditions. In the present work, Baffin Bay samples from both a fixed station (Ice Camp) and an ice-breaker cruise (Amundsen) were sampled for phytoplankton isolation before, during and at the peak of the Arctic spring bloom. More than 1,000 cultures were obtained by serial dilution, single cell pipetting and flow cytometry (FCM) cell sorting, characterized by partial 18S rRNA sequencing and optical microscopy and de-replicated to a subset of 276 strains presented here.

## Material and Methods

### Sampling

The Green Edge project (http://www.greenedgeproject.info) aimed at investigating the dynamics of the Arctic spring bloom at the ice edge. Samples for phytoplankton isolation were obtained both at a fixed station (Ice Camp) and during a cruise on-board the Canadian ice breaker CCGS Amundsen.

The Ice Camp (IC) was set up near the Inuit village of Qikiqtarjuaq, Nunavut, in Baffin Island (67*^◦^* 28’ N, 63*^◦^* 47’ W), in a location identified to have little influence from continental drainage (Figure 1). To observe the changes in the phytoplankton community during the ice melting process, sampling was carried out between May 4^th^ and July 18^th^ 2016. Samples were collected in the water column under the ice at two depths three times a week and from melted ice cores once per week. The ice cores were melted at room temperature with the addition of 0.2 *µ*m filtered sea water prior to isolation procedures.

**Figure 1.**
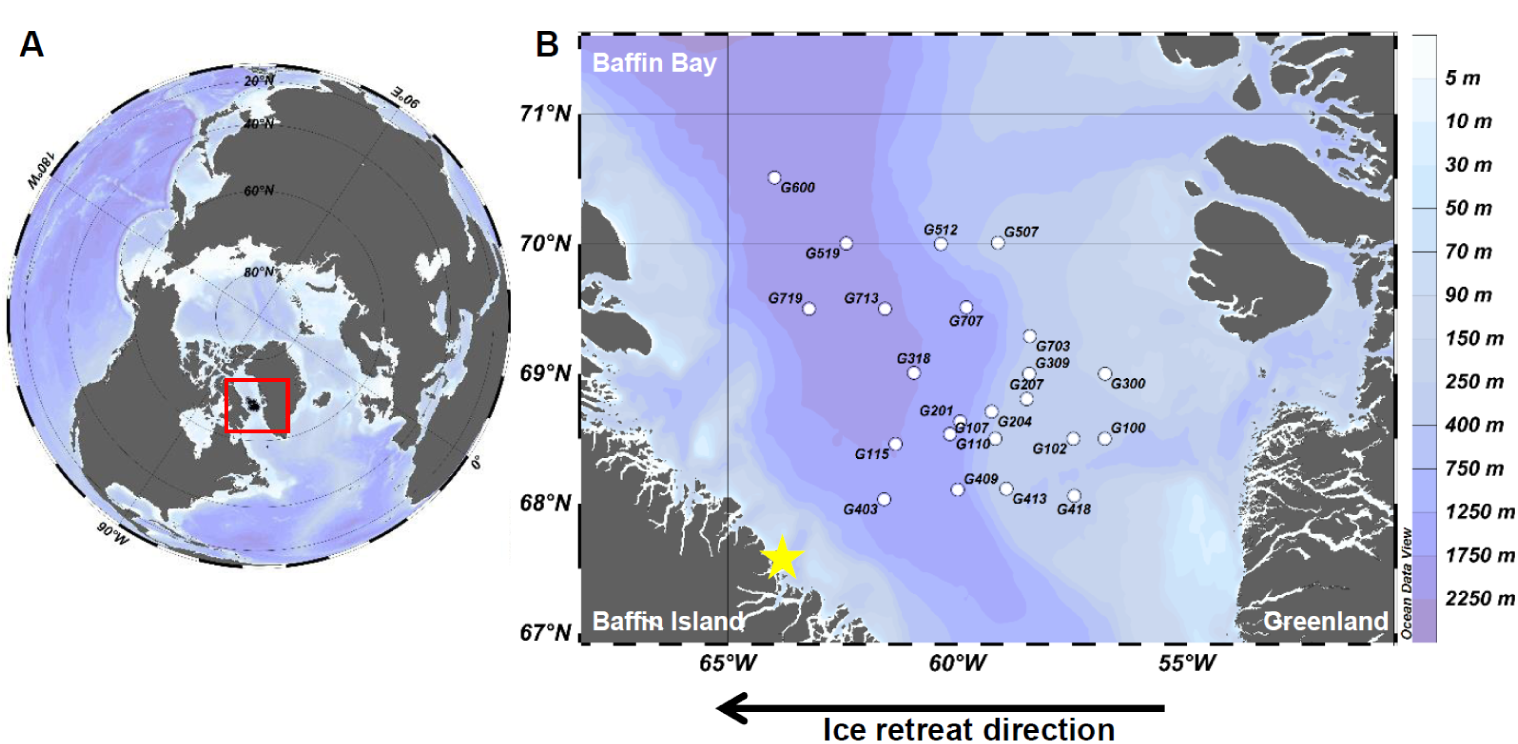
Sampling stations. Sampling stations where phytoplankton strains were retrieved. A) Sampling region (red square). B) The yellow star indicates the location of the Green Edge Ice Camp (IC) (67.48 *^◦^* N,-63.79 *^◦^* W); Amundsen (AM) cruise stations are marked by white dots; black arrow indicates the ice retreat direction during the melting process.

The Amundsen cruise (AM) took place between June 3^rd^ and July 14^th^ 2016 in Baffin Bay, Canada, between 60*^◦^* N and 70*^◦^* N. (Figure 1). Sampling transects were designed to cross the Marginal Ice Zone perpendicularly in order to observe changes in the phytoplankton community from open water to solid sea ice (Figure 1). Seawater for isolation was sampled approximately every two days at two depths with Niskin bottles mounted on a CTD frame Sea-Bird SBE-911 plus.

The development of the spring phytoplankton bloom at the Ice Camp site was monitored by flow cytometry (Lopes dos Santos et al. *in prep*) and its phases were defined as follows: ‘pre-bloom’ from May 4 to May 23; ‘bloom-development’ from May 24 to June 22 and ‘bloom-peak’ from June 23 to July 18. Amundsen strains were not related to bloom phases due to spatial variability across the Marginal Ice Zone during sampling.

### Strain isolation and maintenance

Several isolation strategies were employed in order to maximize the number and diversity of cultures retrieved. Different pre-isolation procedures were applied to different samples, which included filtration, concentration and enrichment. In order to target the smaller plankton size fractions, samples were gravity pre-filtered with 3 *µ*m and 0.8 *µ*m filters prior to enrichment or serial dilution, as described previously (Balzano et al., 2017; Le Gall et al., 2008). Some samples were concentrated with tangential flow filtration (Vivaflow Cartridge 200, Sartorius) with a 0.2 *µ*m polyethersulfone membrane using 2 L of seawater or 0.5 L of melted ice core. Enrichment was performed by mixing 25 mL of pre-filtered seawater with 1 mL of L1 (Guillard and Hargraves, 1993) or PCR-S11 culture medium (Rippka et al., 2000) (media recipes at http://roscoff-culture-collection.org/protocols/media-recipes). Diatom proliferation was prevented in some cultures by the addition of GeO_2_ (Sigma-Aldrich, St-Quentin-Fallavier, France) at 9.6*µ*M.

Isolation from enriched samples was performed by single cell pipetting or by FCM cell sorting using a FACSAria cytometer (Becton Dickinson, San José, CA, USA). In order to prevent cell damage, cells were sorted in K medium (Keller et al., 1987) with 0.01% BSA concentration as described previously (Marie et al., 2017).

For serial dilution either 500 or 50 *µ*L of water sample was added to 15 mL of L1. Then, 24 wells of a Greiner Bio-One^TM^ 96 Deep Well plate (Dominique Dustscher, Brumath, France) were filled with 0.5 mL of each dilution. Wells were later screened by optical microscopy and with a Guava*Q*_R_(Merck, Darm-stadt, Germany) flow cytometer. Unialgal wells were transferred to ventilated T-25 CytoOne*Q*_R_flasks (Starlab, Orsay, France) with 15 mL of L1 media.

All cultures were incubated at 4*^◦^* C with a 12:12 light–dark cycle and transferred to new medium once a month. Light intensity was approximately 100 *µ*mole photons.m^-2^.s^-1^. The isolation method, culture medium and environmental conditions for each strain are reported in Supplementary Data S1.

Cultures were screened and de-replicated by optical microscopy and partial 18S rRNA sequences (see below). We aimed to keep, whenever possible, one strain of each taxon per sampling day and per depth. After de-replication, 416 strains were added to the Roscoff Culture Collection (http://www.roscoff-culture-collection.org) of which 276 were chosen to be described in this paper based on 18S rRNA sequence quality and reliability of culture growth.

### Molecular analyses

Strains were identified using the V4 region of the 18S rRNA gene. DNA was extracted directly from the cultures by a simple heating cycle of 95*^◦^*C for five minutes, prior to PCR. A DNA extraction with NucleoSpin Plant II kit (Macherey-Nagel) was performed following the manufacturer’s instructions for thick-walled or low concentration strains. For 18S rRNA amplification the primers 63F (5’-ACGCTT-GTC-TCA-AAG-ATTA-3’) and 1818R (5’-ACG-GAAACC-TTG-TTA-CGA-3’) (Lepère et al., 2011) were used. PCR amplification was per-formed in a 10 *µ*L mix containing 5 *µ*L of Phusion High-Fidelity PCR Master Mix*Q*_R_2*×*, 0.3 *µ*M final concentration of primer 63F, 0.3 *µ*M final concentration of primer 1818R, 1 *µ*M of DNA and H_2_O. Thermal conditions were: 98 *^◦^* C for 5 min, followed by 35 cycles of 98 *^◦^* C for 20 s, 55 *^◦^* C for 30 s, 72 *^◦^* C for 90 s, and a final cycle of 72 *^◦^* C for 5 min. For most of the cultures the 18 S rRNA gene was sequenced using the internal primer 528F (5’-CCG-CGG-TAATTC-CAG-CTC-3’) (Zhu et al., 2005).

Partial sequences were compared with those available in Genbank using the BLAST plugin in Geneious 10 (Kearse et al., 2012). Sequences were aligned using the ClustalW plugin in Geneious 10 and grouped into genotypes with 100% sequence similarity. Genotypes represented by more than one strain are listed in Supplementary Data S1. Phylogenetic trees were built using the maximum likeli-hood (ML) method with bootstrap values estimated with 1,000 replicates (Felsen-stein, 1985) using PhyML (Guindon et al., 2010) as implemented in Geneious 10.

### Microscopy

One strain per genotype representative of the 18S rRNA genetic diversity was cho-sen for optical light microscopy (LM). Using a Nikon Eclipse 80i (Nikon) with a 100x objective and differential interference contrast, pictures of live cultures were captured with a SPOT digital camera (Diagnostics Instruments, Sterling Heights, MI, USA).

## Results

In the present study, 18S rRNA gene sequences and light microscopy were used to characterize 276 Arctic strains obtained during the Green Edge campaign (Supplementary Data S1), 77 and 199 isolated from ice and water samples, respectively (Figure 2). By combining different pre-isolation and isolation techniques we were able to retrieve 276 strains assigned to 57 genotypes characterized by 100% similarity of partial 18S rRNA sequencing. There was a significant level of novelty within these strains since almost 60% of the representative sequences of genotypes did not match any sequence from previously cultured strains (Table 1) and more than 40% did not match any existing sequence from environmental datasets. The sequence of one strain (**RCC5319**) had only a 95.3 % match to any existing sequence. Strains belonged to 5 divisions (Table 2): Heterokontophyta (208), Chlorophyta (44), Cryptophyta (16), Haptophyta (4) and Dinophyta (4). Diatoms were by far the most abundant group (193) with the genera *Chaetoceros* (42) and *Attheya* (40), followed by *Synedra* (23), *Thalassiosira* (18), Naviculales (16) and *Fragilariopsis* (17) being the most represented. The flagellates *Rhodomonas* (16) and *Pyramimonas* (24) were the most abundant non-diatom genera. With 10 strains, *M. polaris* dominated picoplanktonic isolates, although one strain of *B. prasinos* was also isolated. Four strains of dinoflagellates assigned to *Biecheleria* sp. were retrieved from samples from the Amundsen cruise. The level of novelty varied between the different taxonomic groups and for some classes such as Chlorophyceae and Cryptophyceae, we did not recover any strains corresponding to novel 18S rRNA sequences (Figure S1).

**Figure 2.**
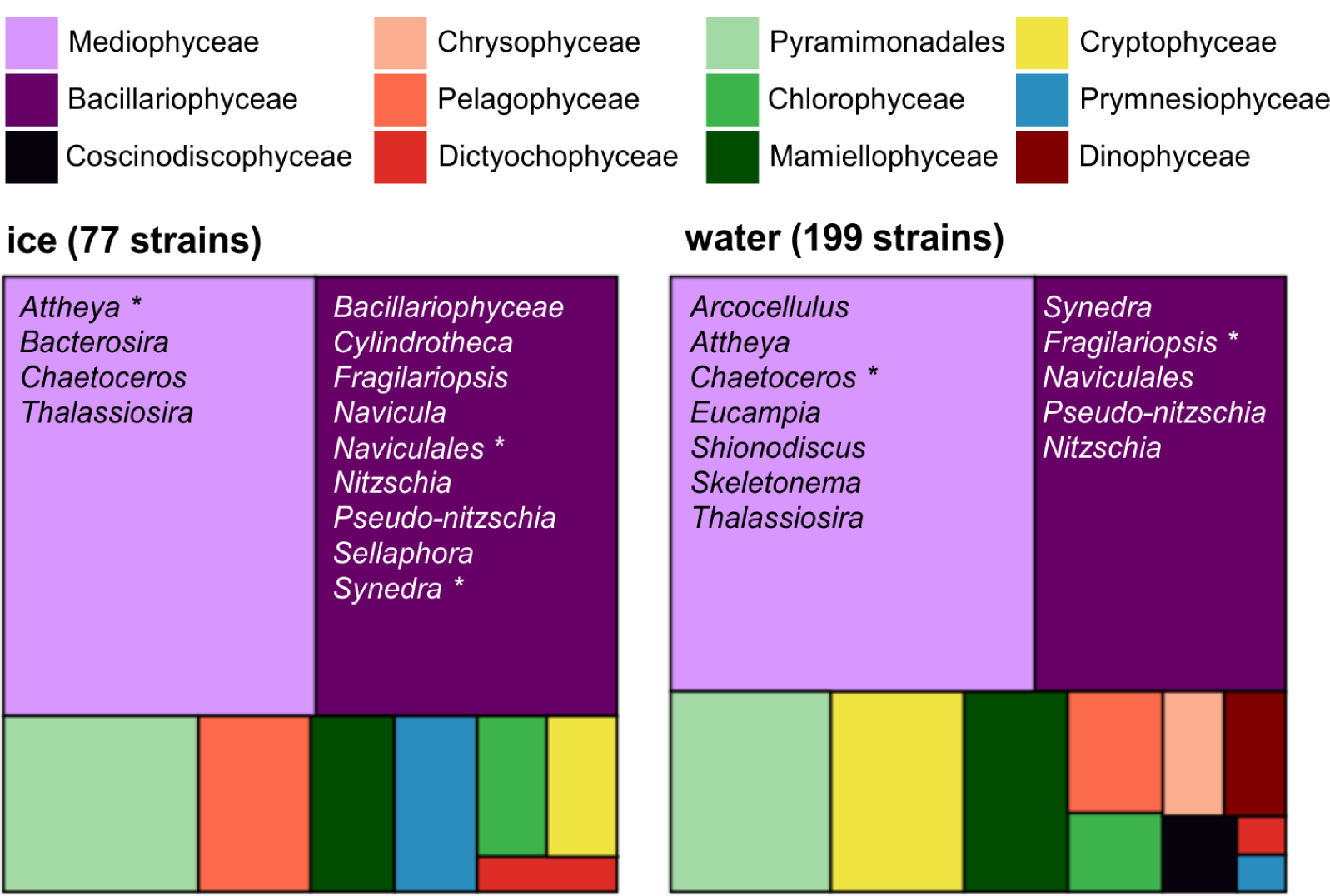
Overall diversity of strains. Overall diversity of the strains retrieved from ice and water samples assigned at the class level. Diatoms genera and most abundant strains are marked with as asterisk.

**Table 1.** Level of novelty of the different genotypes based on BLAST analysis of 18S rRNA against Genbank nr database (Supplementary Data S2).

| Genotype novelty<sup>1</sup> | Number of genotypes |
| --- | --- |
| cultured | 24 |
| detected - uncultured | 9 |
| undetected | 24 |

<sup>1</sup> "cultured" corresponds to genotypes which 18S rRNA sequence is 100% similar to that of a culture that has been isolated previously, "detected - uncultured" correspond to genotypes which 18S rRNA sequence is 100% similar to that of a sequence detected in the environment but for which no culture existed prior to this work and "undetected" corresponds to genotypes for which no 100 % similar 18S rRNA sequence had been detected previously in the environment.

**Table 2.** Number of strains obtained from water and ice samples for each genus.

| Division | Genus | water | ice |
| --- | --- | --- | --- |
| Chlorophyta | *Bathycoccus* |  | 1 |
|  | *Chlamydomonas* | 4 | 2 |
|  | *Mantoniella* | 2 | 1 |
|  | *Micromonas* | 9 | 1 |
|  | *Pyramimonas* | 17 | 7 |
| Cryptophyta | *Rhodomonas* | 14 | 2 |
| Dinophyta | *Biecheleria* | 4 |  |
| Haptophyta | *Isochrysis* |  | 3 |
|  | *Pseudohaptolina* | 1 |  |
| Heterokontophyta | *Actinocyclus* | 2 |  |
|  | *Arcocellulus* | 3 |  |
|  | *Attheya* | 27 | 13 |
|  | *Bacterosira* |  | 1 |
|  | *Chaetoceros* | 36 | 6 |
|  | *Coscinodiscus* | 1 |  |
|  | *Cylindrotheca* |  | 3 |
|  | *Dinobryon* | 1 |  |
|  | *Eucampia* | 1 |  |
|  | *Fragilariopsis* | 14 | 3 |
|  | *Navicula* |  | 2 |
|  | Naviculales | 10 | 6 |
|  | *Nitzschia* | 4 | 3 |
|  | Pedinellales |  | 1 |
|  | Pelagophyceae | 6 | 4 |
|  | *Pseudo nitzschia* | 8 | 1 |
|  | *Sellaphora* |  | 1 |
|  | *Shionodiscus* | 2 |  |
|  | *Skeletonema* | 1 |  |
|  | *Spumella* | 3 |  |
|  | *Stauroneis* |  | 2 |
|  | *Synedra* | 18 | 5 |
|  | *Synedropsis* | 1 | 1 |
|  | *Thalassiosira* | 10 | 8 |

### Phylogenetic analysis of culture diversity

#### Diatoms - Bacillariophyceae

The *Cylindrotheca* sp. genotype represented by **RCC5463** contains two strains from ice core samples from the pre-bloom and bloom-development phases (Supplementary Data S1). Cells are solitary with two chloroplasts, a long apical (> 35 *µ*m) and short transapical axis (~ 3 *µ*m) (Figure 3R). *Cylindrotheca* is a genus frequently observed in the Arctic, mainly represented by the cosmopolitan species complex *C. closterium* (Katsuki et al., 2009; Lovejoy et al., 2002; Poulin et al., 2011; Li et al., 2007). However, sequences from the strains obtained in this study branched apart from *C. closterium* (Figure 4), but grouped with 100% identity with an uncultured *Cylindrotheca* sequence from the Arctic (GenBank JF698839).

**Figure 3.**
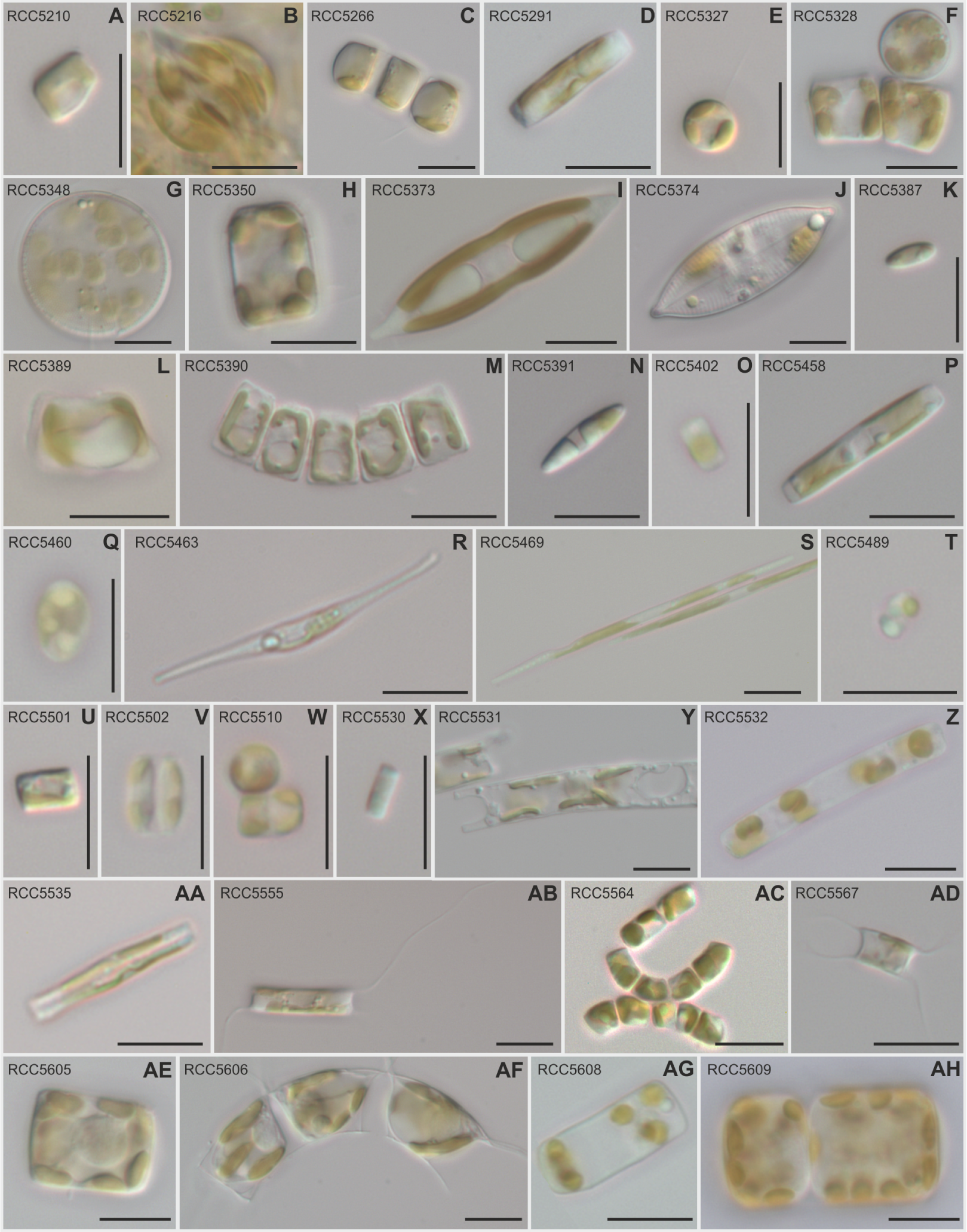
Light microscopy images of diatom strains - see legend on next page. Light microscopy images from diatoms strains retrieved during Green Edge 2016 campaign. Size bars correspond to 10 *µ*m. **A**) *Chaetoceros neogracilis* strain RCC5210; **B**) Cylindrotheca sp. strain RCC5216; **C**) *Chaetoceros gelidus* strain RCC5266 forming a small chain; **D**) *Synedropsis hyperborea* strain RCC5291; **E**) *Tha-lassiosira* sp. strain RCC5327; **F**) *Bacterosira bathyomphala* strain RCC5328 in both girdle and valve view; **G**) *Thalassiosira* cf *antarctica* strain RCC5348 valve with fine radiating areolae; **H**) *Thalassiosira* sp. strain RCC5350 in girdle view; **I**) *Navicula* sp. strain RCC5373; **J**) *Navicula ramosissima* strain RCC5374; **K**) Naviculales sp. RCC5387 cell in valve view; **L**) *Nitzschia* sp. strain RCC5389; **M**) *Nitzschia* sp. RCC5390 cells in ribbon-like colonies; **N**) *Nitzschia* sp. strain RCC5391; **O**) Bacillariophyceae strain RCC5402; **P**) *Nitzschia* sp. strain RCC5458; **Q**) *Sellaphora* sp. strain RCC5460; **R**) *Cylin-drotheca* sp. strain RCC5463; **S**) *Pseudo-nitzschia arctica* strain RCC5469; **T**) *Nitzschia* sp. strain RCC5489; **U**) *Fragilariopsis cylindrus* strain RCC5501; **V**) *Skeletonema* sp. strain RCC5502; **W**) *Nitzschia* sp. strain RCC5510; **X**) *Arcocellulus* sp. strain RCC5530; **Y**) *Eucampia groenlandica* strain RCC5531; **Z**) *Shionodiscus bioculatus* strain RCC5532; **AA**) *Synedra* sp. strain RCC5535; **AB**) *Attheya longicornis* strain RCC5555 solitary cell in girdle view; **AC**) Naviculales sp. RCC5564 forming a small chain; **AD**) *Attheya septen-trionalis* strain RCC5567; **AE**) *Thalassiosira rotula* RCC5605; **AF**) *Chaetoceros decipiens* strain RCC5606 forming a small, curved chain; **AG**) *Actinocyclus* sp. RCC5608; **AH**) *Shionodiscus bioculatus* strain RCC5609.

**Figure 4.**
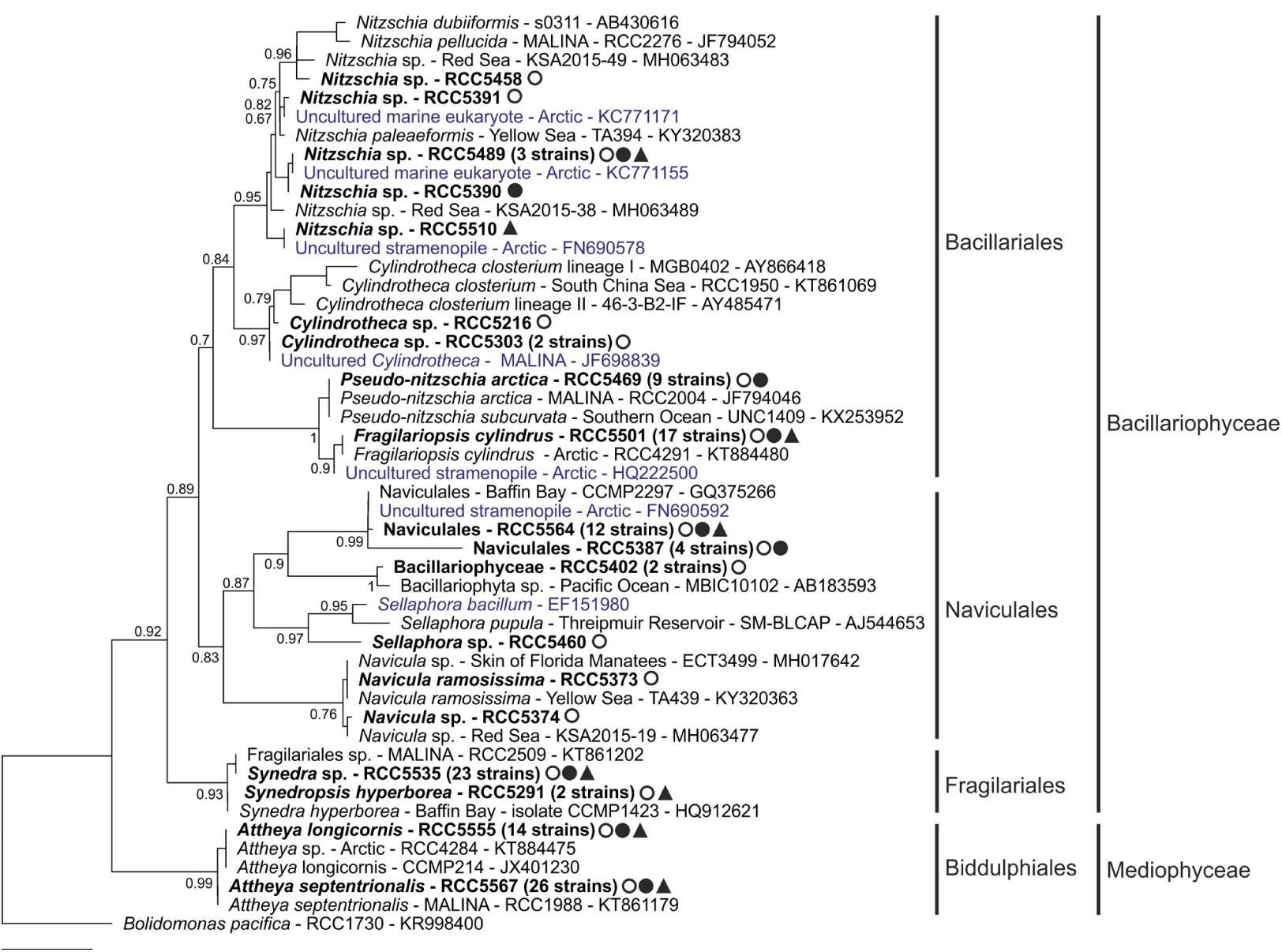
18S rRNA phylogenetic tree of pennate diatoms. 18S rRNA phylogenetic tree inferred by maximum likelihood (ML) analysis for pennate diatom strains obtained during the Green Edge campaign (in bold), using an alignment of 59 sequences with 406 positions. Circles mark strains retrieved from the Ice Camp ice (open) and water samples (solid); triangles (solid) mark Amundsen cruise water samples. The origin, sampling substrate and phase of the bloom from which they were recovered are provided along with their names and RCC code in Supplementary Data S1. When one genotype is represented by more than one strain, the number of strains is indicated between parenthesis. For the reference sequences, the strain (whenever available) and the Genbank ID number are displayed. Environmental sequences are marked in blue.

The sequence of *Cylindrotheca* sp. **RCC5216**, also isolated from an IC ice core sample (Supplementary Data S1), differed from that of *Cylindrotheca* sp. **RCC5463** by two base pairs. Cells of **RCC5216** are curved to sigmoid forming coarse aggregates, with 17-20 *µ*m apical and *∼* 4 *µ*m transpical axes (Figure 3B).

The *Fragilariopsis cylindrus* genotype represented by **RCC5501** groups 17 strains originating from all main sampling sites, substrates and phases of the bloom (Supplementary Data S1). The 18S rRNA sequence matched with 100% similarity the Arctic *F. cylindrus* strain RCC4291 (Figure 4), a known cold-adapted diatom (Mock et al., 2017), used as an indicator of polar water and ice (Quillfeldt, 2004). Cells have a short apical axis (~4 *µ*m), rounded ends, and a transapical axis of approximately 3 *µ*m length (Figure 3U).

*Navicula ramosissima* strain **RCC5373** was retrieved from an ice core sample from the pre-bloom period and shared 100% similarity with *Navicula ramosissima* strain TA439 from the Yellow Sea and *Navicula* sp. strain ECT3499 obtained from the skin of Florida manatees (Figure 4). Cells are solitary, lanceolate, with apical and transpical axes of ~ 25 *µ*m and 7 *µ*m, respectively, two elongated chloroplasts on each side of the girdle, and large lipid bodies (Figure 3I). Interestingly, none of the *Navicula* sp. strains recovered in this study were related to previous polar strains or environmental sequences, despite this genus being diverse (Katsuki et al., 2009) and abundant in the Arctic (Kauko et al., 2018; Poulin et al., 2011).

*Navicula* sp. strain **RCC5374** was recovered from an ice core sample from the bloom-development phase. The sequence of this strain is not very closely related to those of previously reported polar *Navicula*, but is 99.2% similar to strain RCC5373 and 99.7% similar to strain KSA2015-19 from the Red Sea (Figure 4). Cells have a ~ 25 *µ*m apical axis, slightly radiating valvar striae and rostrate ends (Figure 3J).

The Naviculales genotype represented by **RCC5564** contains 12 strains from all phases and sampling sites (Supplementary Data S1). Its sequence is 99.7% similar to Naviculales strain CCMP2297 from northern Baffin Bay and to uncultured sequences from the Arctic (Figure 4). Cells have ~ 3 *µ*m apical and 5 *µ*m pervalvar axes. They are solitary or form short chains (Figure 3AC).

The Naviculales genotype represented by **RCC5387** contains four strains from IC water and ice samples (Supplementary Data S1). Its sequence has low similarity to sequences from GenBank or to the genotype represented by RCC5564, sharing only 96.9% similarity with strain CCMP2297 (Naviculales) (Figure 4). Cells are elongated, mainly solitary, with up to 6 *µ*m apical and 3 *µ*m pervalvar axes (Figure 3K).

The *Nitzschia* sp. genotype represented by **RCC5489** contains three strains from both sampling sites and substrates (Supplementary Data S1). Its sequence has no close similarity to any GenBank sequence from strains besides **RCC5390** (Figure 4). Cells are ~ 11 *µ*m wide, mainly solitary or forming small aggregates (Figure 3L). Members of the genus *Nitzschia* are often reported to thrive in the Arctic (Johnsen et al., 2018; Kauko et al., 2018), *Nitzschia frigida*, for example, being considered as the single most important diatom in association with sea ice (Leu et al., 2015). Surprisingly, none of the *Nitzschia* sp. strains isolated in this study had high 18S rRNA similarity to other known polar strains. They did, however, have high similarity with Arctic environmental sequences (Figure 4).

*Nitzschia* sp. **RCC5390** was retrieved from an IC pre-bloom sample (Supplementary Data S1) and its sequence is the closest to **RCC5489** (99.8% similarity, Figure 4). Cells have 7 *µ*m apical axis and 5 *µ*m pervalvar distance, forming ribbon-like colonies (Figure 3M).

*Nitzschia* sp. **RCC5391** was isolated from an ice core sample during the prebloom period. Its sequence matches with only 97.8% similarity that of a strain TA394 (*Nitzschia paleaeformis*) from the Yellow Sea (Figure 4). Cells are solitary lanceolate with bluntly rounded apices, measuring ~ 10 *µ*m and 2 *µ*m for the apical and transapical axes, respectively (Figure 3N).

*Nitzschia* sp. **RCC5458** was also retrieved from an ice sample from the prebloom period and its sequence is 98.1% similar to *Nitzschia* sp. strain KSA2015-49 from the Red Sea (Figure 4). Cells are linear to lanceolate and larger than other *Nitzschia* strains retrieved in this study, with an apical axis up to 15 *µ*m (Figure 3P).

*Nitzschia* sp. **RCC5510** was isolated from AM waters (Supplementary Data S1). Its sequence is 98.6% similar to *Nitzschia* sp. strain KSA2015-38 from the Red Sea (Figure 4). It is the only strain from this genus recovered only from AM. Its sequence branches apart from all other *Nitzschia* sp. (Figure 4). Cells are almost round in the valvar view and rather small (apical axis ~ 4 *µ*m) compared to the other *Nitzschia* strains isolated here (Figure 3W).

The *Pseudo-nitzschia arctica* genotype represented by **RCC5469** contains nine strains of the recently described *P. arctica* (Percopo et al., 2016), all originating from IC (Supplementary Data S1). Their sequence is 100% similar to *P. arctica* RCC2004 (Figure 4), a potentially endemic species with a wide distribution in the Arctic (Balzano et al., 2017; Percopo et al., 2016). Only solitary cells were observed, with lanceolate shape in valvar view, measuring ~ 50 *µ*m and 3 *µ*m for the apical and transapical axes, respectively (Figure 3S).

The Bacillariophyceae genotype represented by **RCC5402** has two strains, both retrieved from IC ice cores during the pre-bloom period (Supplementary Data S1), and could not be assigned to any specific genus. Their sequence shares 99.2% identity with the Bacillariophyta MBIC10102 strain from the Pacific Ocean and groups with Naviculales sequences with moderate bootstrap support (91%) (Figure 4). Cells are small ~ 4 *µ*m long and 2 *µ*m wide, sometimes solitary, but mainly forming large aggregates (Figure 3O).

*Sellaphora* sp. strain **RCC5460** was retrieved during pre-bloom from an IC ice core sample. Its sequence matches with 99% similarity that of the freshwater *Sellaphora pupula* strain SM-BLCAP (Figure 4). Cells are small, with 5*µ*m apical and 4 *µ*m pervalvar axes, solitary or forming aggregates (Figure 3Q). *S. pupula* is a species complex containing many pseudo- and semi-cryptic representatives capable of thriving is a wide range of environmental conditions (Poulíčková et al., 2008). Further molecular/morphological analyses are needed to properly assign this genotype.

The *Synedropsis hyperborea* genotype represented by **RCC5291** contains only two strains, from both IC and AM (Supplementary Data S1). Its sequence shares 100% similarity with *S. hyperborea* strain CCMP1423 (Figure 4), although members of the Fragilariaceae are not well resolved by 18S rRNA (Balzano et al., 2017). Cells are solitary or in pairs, exhibiting great variability in shape, which is attributed to vegetative cell division (Hasle et al., 1994). The apical axis is ~ 14 *µ*m (Figure 3D). *S. hyperborea* is an Arctic species with circumpolar distribution, often found in association with sea ice and as an epiphyte of *Melosira arctica* (Assmy et al., 2013; Hasle et al., 1994).

The *Synedra* sp. genotype represented by **RCC5535** comprises ten strains of which four were isolated from the Amundsen cruise and the other six from within or under the IC ice (Figure 4). Its sequence shares 100% identity with other Arctic strains such as Fragilariales RCC2509. Cells vary considerably in shape, from almost linear to lanceolate and sometimes asymmetrical in the valvar central area. Apical and transapical axes are *∼* 13 *µ*m and 3 *µ*m, respectively (Figure 3AA).

#### Diatoms - Coscinodiscophyceae

The *Actinocyclus* sp. genotype represented by **RCC5608** comprises two strains isolated from AM waters during the bloom-peak (Supplementary Data S1). Its sequence shares 100% similarity with a clone from the Arctic (EU371328), and 99.8% with the *Actinocyclus* sp. MPA-2013 isolate from the Pacific Ocean (Figure 5). Cells have a pervalvar axis (13-17 *µ*m) longer than the valvar diameter (~ 5 *µ*m) and discoid chloroplasts (Figure 3AG). Although sometimes spotted in low abundance (Crawford et al., 2018; Katsuki et al., 2009), this genus may dominate phytoplankton biomass in Arctic spring blooms (Lovejoy et al., 2002).

*Coscinodiscus* sp. strain **RCC5319** was isolated from an IC under-ice sample at the peak of the bloom (Supplementary Data S1). The sequence is only 95% similar to that of *Coscinodiscus jonesianus* isolate 24VI12 (KJ577852) (Figure 5). Unfortunately, this strain was lost and no images are available. *Coscinodiscus* may be abundant under the ice pack (Duerksen et al., 2014) and is often reported in Arctic diversity studies (Booth et al., 2002; Lovejoy et al., 2002).

**Figure 5.**
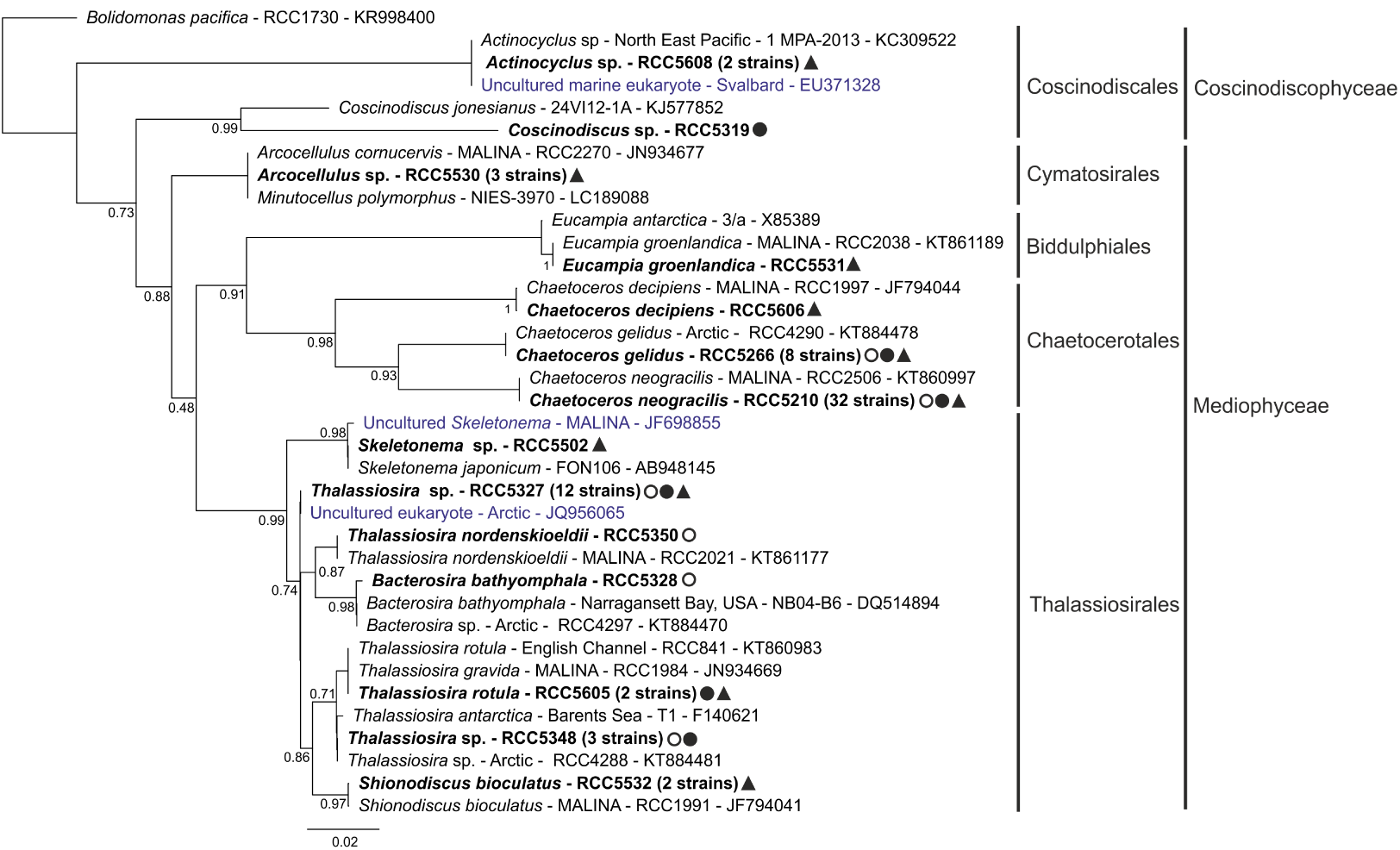
18S rRNA phylogenetic tree of centric diatoms. 18S rRNA phylogenetic tree inferred by maximum likelihood (ML) analysis for centric diatoms strains. Legend is the same as in Figure 4, using an alignment of 55 sequences with 593 positions.

#### Diatoms - Mediophyceae

The *Arcocellulus* sp. genotype represented by **RCC5530** contains three strains isolated from 17 m depth from the Amundsen cruise (Supplementary Data S1). Their sequence is 100% similar to RCC2270 *Arcocellulus cornucervis* (Figure 5). However, 18S rRNA sequences do not have enough resolution to separate *Arcocellulus* sp. from closer groups such as *Minutocellulus* sp. (Balzano et al., 2017), requiring further analyses for proper assignation. Cells are small (~ 5 *µ*m) and solitary (Figure 3X). The cold adapted *A. cornucervis* has been reported to be part of the protist community in the Arctic (Blais et al., 2017), including in Baffin Bay in early summer (Lovejoy et al., 2002).

The *Attheya septentrionalis* genotype represented by **RCC5567** comprises 26 strains from all substrates and sampling sites, from bloom-development and bloom-peak phases (Supplementary Data S1). Their sequence shares 100% similarity with the Arctic strain RCC1988 (Figure 5). Cells are lightly silicified with ~ 6 *µ*m pervalvar axis and horns up to two times the cell length. They are either solitary or form big aggregates (Figure 3AD). *A. septentrionalis* is often reported in abundance in Arctic waters and ice (Assmy et al., 2013; Balzano et al., 2017), outcompeting pennate diatoms in high-luminosity/low nutrient conditions (Campbell et al., 2017).

The *Attheya longicornis* genotype represented by **RCC5555** contains 14 strains, 11 of which were retrieved from Amundsen water samples (Supplementary Data S1). Sequences are 100% identical to the Arctic *A. longicornis* strains RCC4284 and CCMP214 (Figure 5). Cells are often solitary or in short chains, with horns up to three times the length of the pervalvar axis (Figure 3AB). Together with *A. septentrionalis*, *A. longicornis* can comprise a significant portion of the diatom community in Arctic sea ice (Campbell et al., 2017).

*Bacterosira bathyomphala* strain **RCC5328** was retrieved from an ice core sample and its sequence shares 99.8% identity with the Arctic *Bacterosira* sp. RCC4297 and with *B. bathyomphala* strain NB04-B6 from an estuary (Figure 5). Cells (~ 9 *µ*m pervalvar axis) form short and tight chains with contiguous valves (Figure 3F). *B. bathyomphala* is often reported in northern and polar waters (Crawford et al., 2018; Johnsen et al., 2018), especially where silicate concentration is high (Luddington et al., 2016).

The *Chaetoceros neogracilis* genotype represented by **RCC5210** contains 33 strains retrieved from all sites, substrates and phases of the bloom (Supplementary Data S1). Its sequences share 100% similarity with polar *C. neogracilis* strains (e.g. RCC2506). The 18S rRNA gene does not, however, have enough resolution to differentiate within *C. neogracilis* clades (Balzano et al., 2017). Cells are small, solitary or forming aggregates, with the perivalvar axis slightly longer than the valvar diameter (4 *µ*m) (Figure 3A). The genus *Chaetoceros* is abundant in temperate and polar waters (Lovejoy et al., 2002; Malviya et al., 2016) and *C. neogracilis* dominates the nanophytoplankton community in surface waters in the Beaufort Sea in the summer (Balzano et al., 2012).

The *Chaetoceros gelidus* genotype represented by **RCC5266** contains eight strains from all substrates and sampling sites, but only from the bloom-development and bloom-peak periods (Supplementary Data S1). Their sequences were 100% similar to those of the Arctic strains RCC4290 and RCC1992 (Figure 5). Cells are rectangular (~ 6 *µ*m), forming small, tight chains with narrow apertures and long inner setae, up to 25 *µ*m (Figure 3C). *C. gelidus* is a recently described species, previously considered as an *Chaetoceros socialis* ecotype, and is characteristic of northern temperate and polar waters (Chamnansinp et al., 2013). It is reported to form blooms (Booth et al., 2002) and can represent an important fraction of diatom abundance and biomass in Baffin Bay (Crawford et al., 2018).

*Chaetoceros decipiens* strain **RCC5606** was isolated from 30 meters depth in AM water (Supplementary Data S1). Its sequence is 99.8% similar to Arctic strain *C. decipiens* RCC1997 (Figure 5). Cells (~ 10-30 *µ*m apical axis) have very long inner setae (> 100 *µ*m) and form short, semi-circular colonies (Figure 3AF), which contrasts with previous morphological descriptions of *C. decipiens* (Balzano et al., 2017; Hasle and Syvertsen, 1997), indicating that it might correspond to a new genotype. This cosmopolitan species has frequently been reported in the Arctic, both in ice and open waters (Joo et al., 2012; Lovejoy et al., 2002; Johnsen et al., 2018).

*Eucampia groenlandica* **RCC5531** strain was retrieved from 30 m depth during the Amundsen cruise. Its sequence shares 100% similarity with *E. groenlandica* Arctic strain *R*CC2038 (Figure 5). Cells are lightly silicified with varying sizes, forming straight or moderately curved colonies (Figure 3Y). *E. groenlandica* was first reported in Baffin Bay (Cleve, 1896) although its distribution is not constrained to the Arctic (Lee and Lee, 2012).

The *Shionodiscus bioculatus* genotype represented by **RCC5532** contains two strains isolated from the Amundsen cruise (Supplementary Data S1). Its sequence shares 99.8% similarity with *S. bioculatus* strain RCC1991 from the Beaufort Sea (Figure 5). The morphology of the two strains differs (Figure 3Z and F1): **RCC5532** cells have a longer pervalvar axis (~ 32 *µ*m), shorter valve diameter and fewer discoid chloroplasts in comparison to **RCC5609**. Isolates with identical 18S rRNA may present cryptic diversity based on ITS divergence (Luddington et al., 2016). *S. bioculatus* is reported as dominating the top portion of submerged sea-ice ridges (Fernández-méndez et al., 2018).

*Skeletonema* sp. **RCC5502** strain was retrieved during the Amundsen cruise and its sequence shared 100% similarity with *S. japonicum* from Onagawa Bay and 99.7% with an Arctic environmental sequence (JF698855, Figure 5). Cells are small (5 *µ*m diameter) with a very short pervalvar axis ~ 3 *µ*m, being either solitary or in pairs (Figure 3V. The genus *Skeletonema* has been reported from high latitude, winter samples (Eilertsen and Degerlund, 2010) and *S.* aff. *japonicum* seems to thrive in polar environments with low silicate concentration (Luddington et al., 2016).

The *Thalassiosira* sp. genotype represented by **RCC5327** contains 12 strains from all sampling sites, substrates and phases of the bloom (Supplementary Data S1). The best match to its sequence is from an Arctic environmental sequence (99.5% similarity), branching apart from other *Thalassiosira* clades (Figure 5). It shares 99.2% identity with *T. nordenskioeldii* strain RCC2021. Cells are small (< 8 *µ*m diameter) with a long pervalvar dimension relative to valve size and long (*>* 20 *µ*m) marginal threads (Figure 3E).

The *Thalassiosira* sp. genotype represented by **RCC5348** contains three strains from IC water and ice. Its sequence is 99.8% similar with a *Thalassiosira antarctica var. borealis* isolate from the Barents Sea (Figure 5). Cells are cylindrical with a short pervalvar axis, a 17-22 *µ*m valvar diameter, and contain fine areolae radiating from the valve center (Figure 3G). *T. antarctica* is reported in coastal and ice-edge cold waters (Hasle and Heimdal, 1968) and associated with high-nutrient concentrations (Luddington et al., 2016).

The sequence of *Thalassiosira nordenskioeldii* strain **RCC5350** isolated from an ice core sample is (100%) identical to that of *T. nordenskioeldii* Arctic strain RCC2021 (Figure 5). Cells are cylindrical, either solitary or forming colonies, with a ~ 6 *µ*m valvar diameter and a 10 *µ*m pervalvar axis, with long processes (Figure 3H). *T. nordenskioeldii* is widely distributed in North Atlantic cold, temperate and polar waters (Crawford et al., 2018; Johnsen et al., 2018), often associated with ice (Luddington et al., 2016).

The *Thalassiosira rotula* genotype represented by **RCC5605** contains two strains, one isolated during the Amundsen cruise and one from under-ice at the Ice Camp during the bloom peak (Supplementary Data S1). The sequence from this genotype had 100% similarity with those of *T. rotula* strains from the Arctic and the English Channel, but also with that of *Thalassiosira gravida* (RCC1984) (Figure 5). 18S rRNA is not a good marker to discriminate between *T. rotula*, a known cosmopolitan species (Hasle and Syvertsen, 1997; Whittaker et al., 2012), and the bipolar *T. gravida* (Balzano et al., 2017). Cells are mainly solitary, with a ~ 6 *µ*m valvar diameter and a 10-13 *µ*m pervalvar axis with several long marginal threads (Figure 3AE).

#### Other Heterokontophyta

*Dinobryon faculiferum* strain **RCC5261** was isolated from 1.5 m depth in IC waters from the peak of the bloom (Supplementary Data S1). Its sequence shares 100% similarity to those of other Arctic strains, such as RCC2294 (Figure 6B). Cells are solitary with a ~ 4 *µ*m diameter lorica and long spines (*>* 25 *µ*m) (Figure 7C). *D. faculiferum* is a frequently observed mixotroph in Arctic surface waters (Balzano et al., 2012; Lovejoy et al., 2002) that can be found encysted in the top section of ice cores (Kauko et al., 2018), although it is not restricted to polar environments (Unrein et al., 2010).

**Figure 6.**
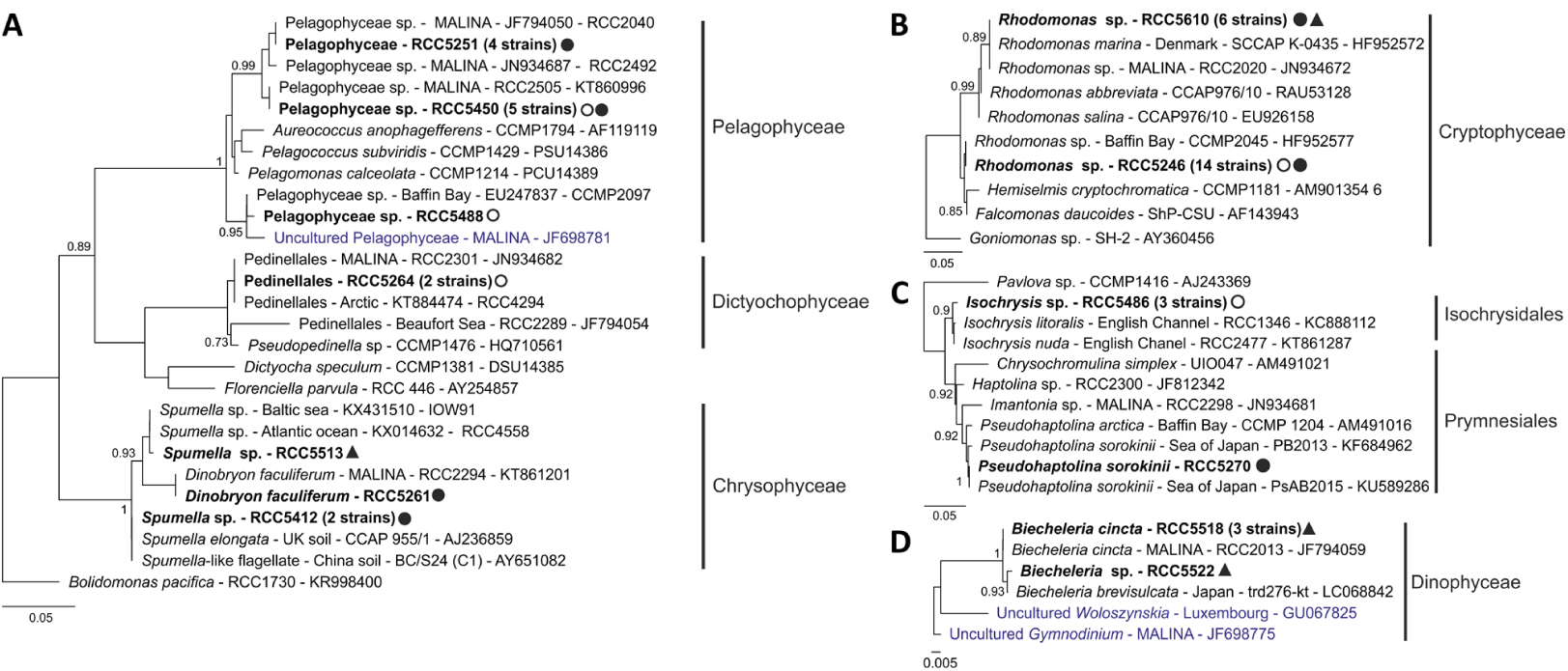
18S rRNA phylogenetic tree of other taxonomic groups. 18S rRNA phylogenetic tree inferred by maximum likelihood (ML) analysis for the strains obtained during the Green Edge campaign (in bold) for: A) Cryptophyta, using an alignment of 15 sequences with 638 positions; B) Heterokontophyta division, alignment of 41 sequences with 396 positions; C) Haptophyta, using an alignment of 15 sequences with 375 positions and D) Dinophyta, alignment of 7 sequences with 300 positions. Legend as in Figure 4.

*Spumella* sp. **RCC5513** strain isolated from an AM sample branches with *D. faculiferum* and its sequence is 99.8 % similar to those of *Spumella* sp. strains from the Baltic Sea (isolate IOW91) and the Atlantic Ocean (RCC4558) (Figure 6B). Cells are colorless and solitary, round or slightly elongated with 4 *µ*m diameter and 5 *µ*m flagella (Figure 7N). Heterotrophic flagellates from the genus *Spumella* have been previously reported in the Arctic (Lovejoy et al., 2006) and are mostly cold-adapted and associated with lower salinities (Grossmann et al., 2015).

**Figure 7.**
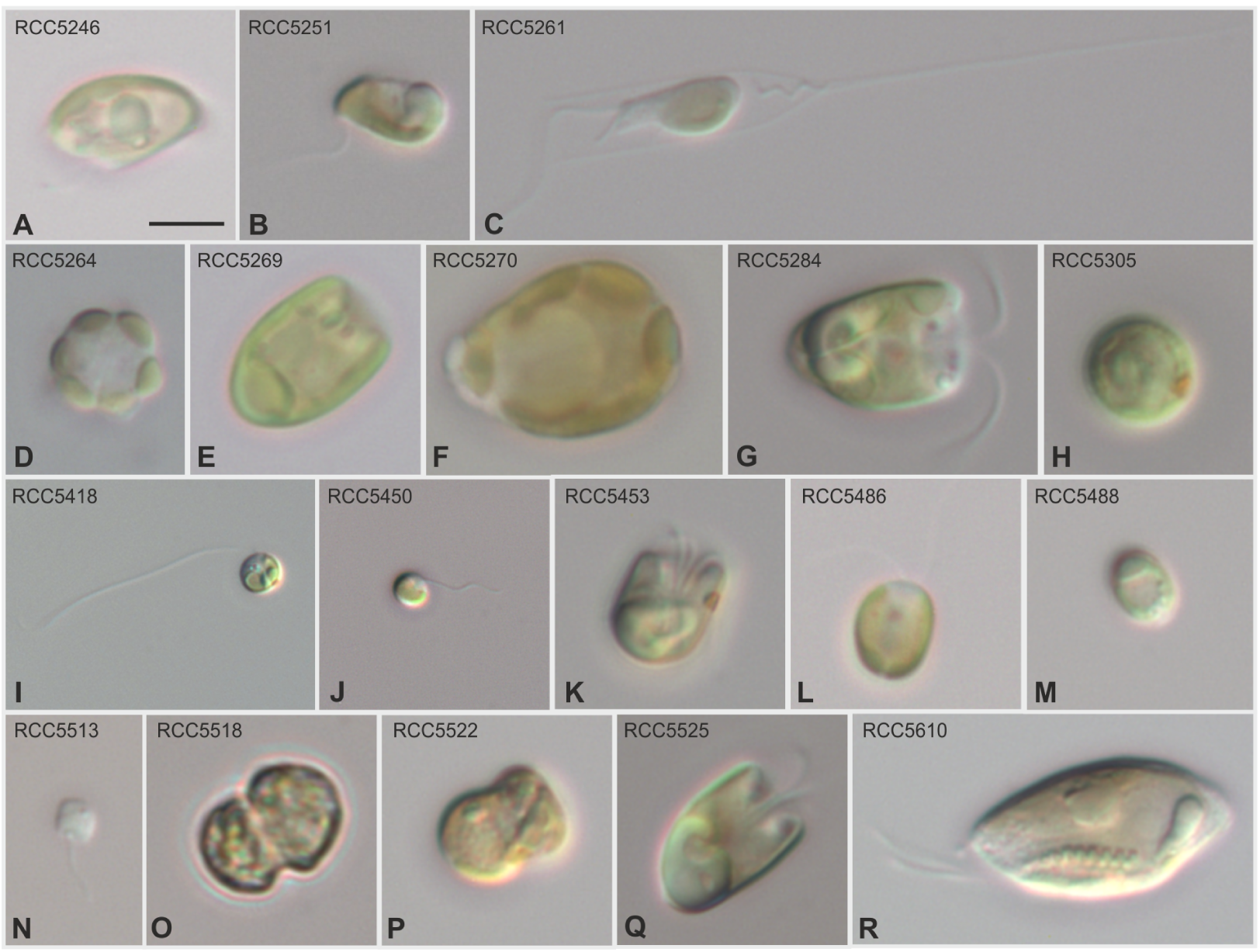
Light microscopy images of flagellate strains. Light microscopy of selected strains of flagellates obtained during Green Edge 2016 campaign. Size bars correspond to 5 *µ* m. **A**) *Rhodomonas* sp. strain RCC5246; **B**) Pelagophyceae strain RCC5251; **C**) *Dinobryon faculiferum* strain RCC5261; **D**) Pedinellales RCC5264 cell showing a ring of six peripheral chloroplasts; **E**) *Pyramimonas australis* strain RCC5269; **F**) *Pseudohaptolina sorokinii* RCC5270; **G**) *Pyramimonas* sp. strain RCC5284; **H**) *Chlamydomonas* sp. strain RCC5305; **I**) *Mantoniella baffinensis* strain RCC5418 with a long flagellum and visible eyespot; **J**) Pelagophyceae strain RCC5450; **K**) *Pyramimonas* sp. strain RCC5453; **L**) *Isochrysis* sp. strain RCC5486; **M**) Pelagophyceae strain RCC5488; **N**) *Spumella* sp. strain RCC5513; **O**) *Biecheleria cincta* strain RCC5518; **P**) *Biecheleria* sp. strain RCC5522; **Q**) *Pyramimonas* sp. strain RCC5525; **R**) *Rhodomonas* sp. RCC5610.

The *Spumella* sp. genotype represented by **RCC5412** contains two isolates from IC waters. Their sequence is 100% similar to those of *Spumella* sp. isolate CCAP 955/1 from a soil sample collected in China and *Spumella elongata* isolate JBC/S24 from the UK (Figure 6B). Interestingly, these sequences are part of a soil sub-cluster within Chrysophyceae clade C with few aquatic representatives (Boenigk et al., 2005). These strains were lost and no images are available.

Pedinellales strain **RCC5264** was retrieved from an IC ice sample at the peak of the bloom (Supplementary Data S1) and its sequence matched with 100% similarity that of the undescribed Pedinellales Arctic strain RCC2301. Cells are solitary, round in anterior view (6 *µ*m diameter), apple-shaped to slightly elongated in side view, with six peripheral chloroplasts (Figure 7D). Further taxonomic analyses are needed to properly assign this strain at the genus level, although its sequence matches with with 98.6 % similarity that of a *Pseudopedinella* sp. strain (CCMP1476) from the Sargasso Sea (Figure 6B).

The Pelagophyceae genotype represented by **RCC5450** groups five strains from IC, four from water samples and one from ice (Supplementary Data S1). Its sequence shares 100% similarity with other Arctic strains such as RCC2505 and RCC2515. Cells are round, *∼* 4 *µ*m in diameter, with two flagella of different size, ~ 2 *µ*m and 7 *µ*m, respectively (Figure 7J). Pelagophyceae may dominate surface waters during the Arctic summer (Balzano et al., 2012) and yet undescribed strains have been recovered previously from northern waters (Balzano et al., 2012).

The Pelagophyceae genotype represented by **RCC5251** contains three strains from the peak of the bloom (Supplementary Data S1) and its representative sequence shares 100% similarity with that of the undescribed Arctic Pelagophyceae RCC2040 (Figure 6B). Cells are elongated with ~ 7 *µ*m in side view (Figure 7B). Pelagophyceae strain **RCC5488**, isolated from an ice sample during bloom-development phase (Supplementary Data S1), has a sequence that branches apart from the other Pelagophyceae genotypes (Figure 6B), matching with 100% similarity another strain isolated from Baffin Bay, CCMP2097. Cells are solitary, ~ 4 *µ*m in size (Figure 7M).

#### Chlorophyta

The *Chlamydomonas* sp. genotype represented by **RCC5305** contains 6 strains isolated from IC water and ice samples from the peak of the bloom and is the only representative of the Chlorophyceae in our set of culture isolates (Supplementary Data S1). Its sequence is 100% identical to sequences from the *Chlamydomonas pulsatilla* polar strain CCCryo 038-99, but also strains from Antarctic ice and Arctic fresh water (Figure 8), all belonging to the *Polytoma* clade (Pocock et al., 2004). Cells are round or elongated, ~ 7 *µ*m in diameter or 10 *µ*m long, respectively (Figure 7H). *Chlamydomonas* is a common genus found in the Arctic during the spring and summer months (Balzano et al., 2012; Lovejoy et al., 2002), that can occur in association with sea-ice (Majaneva et al., 2017).

**Figure 8.**
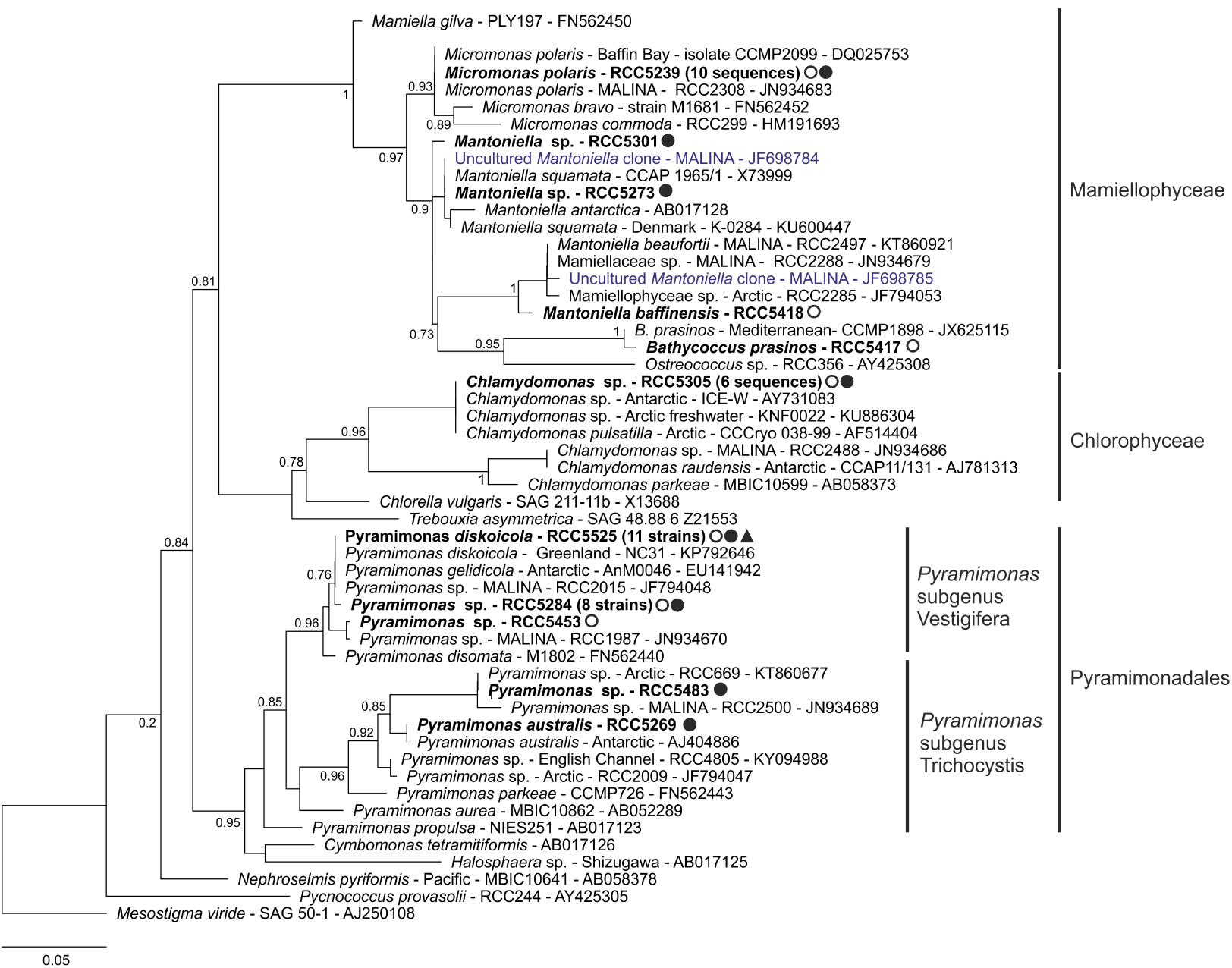
18S rRNA phylogenetic tree of Chlorophyta. 18S rRNA phylogenetic tree inferred by maximum likelihood (ML) analysis for the Chlorophyta strains. Legend is the same as in Figure 4, using an alignment of 70 sequences with 360 positions.

*Bathycoccus prasinos* **RCC5417** strain was recovered from an IC ice core sample during bloom-development (Supplementary Data S1). This genus has recently been observed in Arctic waters (Kilias et al., 2014; Terrado et al., 2013), including during winter (Joli et al., 2017) and has a highly conserved 18S rRNA. Its sequence shares 100% similarity with *Bathycoccus prasinos* strain CCMP1898 from the Mediterranean Sea (Figure 8).

The *Micromonas polaris* genotype represented by **RCC5239** regroups ten strains recovered from Ice Camp ice and water samples. Its sequence shares 100% similarity with those of the Arctic strains *M. polaris* CCMP2099 and RCC2308 (Figure 8). *M. polaris* often dominates the picoplanktonic community in the Arctic (Not et al., 2005; Sherr et al., 2003; Balzano et al., 2012) and metagenomic data suggest its presence in Antarctic waters (Delmont et al., 2015; Simmons et al., 2015).

*Mantoniella baffinensis* **RCC5418**, recently described (Yau et al., 2018), was recovered from pre-bloom IC ice core samples. Its sequence branched apart from other known strains (Figure 8), matching with 98% similarity the Arctic strains RCC2497 and RCC2288 which were also recently described as *Mantoniella beau-fortii* (Yau et al., 2018). Cells are round, ~ 5 *µ*m in diameter bearing two unequal flagella with a visible red eyespot opposite to the flagella (Figure 7I).

*Mantoniella* sp. strain **RCC5273** was isolated from a sample taken at 20 m depth during the peak of the bloom. Its sequence shared 99.8% similarity with that of *Mantoniella squamata* strain CCAP 1965/1, a cosmopolitan species (Hasle and Syvertsen, 1997) frequently observed in the Arctic (Lovejoy et al., 2007; Ma-janeva et al., 2017). This strain was lost and no images are available.

*Mantoniella* sp. strain **RCC5301** was also isolated from 20 m depth during the peak of the bloom and its sequence is not closely related to any strain or environmental sequence. However, it clustered together with other *Mantoniella* sequences, sharing 98.3% identity with *M. squamata* CCAP 1965/1 (Figure 8). This strain was also lost and no images are available.

The *Pyramimonas diskoicola* / *Pyramimonas gelidicola* genotype represented by **RCC5525** contains 11 strains from all main sampling sites, substrates and phases of the bloom (Supplementary Data S1). The sequence from **RCC5525** is 100% similar to that of the Arctic *P diskoicola* and the Antarctic *P. gelidicola* within the subgenus *Vestigifera* (Figure 8). Three types of cell morphology have been observed: pyramidal, elongated and nearly round. A big starch grain with two lobes surrounds a pyrenoid located at the basal end; large lipid bodies are present near the apical end. Cells are ~ 7 *µ*m in length and have four flagella with similar size (Figure 7Q).

The *Pyramimonas sp.* genotype represented by **RCC5284** contains 8 strains from the IC during the later phases of the bloom, 7 of which were isolated from water samples (Supplementary Data S1). The representative sequence shares 99.7% similarity with that of *P. diskoicola* **RCC5525** (Figure 8). Cells are pyra-midal to round, ~ 8 *µ*m long with a pyrenoid and basally positioned starch grain, four flagella shorter than cell length, and a flagellar pit ~ 2 *µ*m deep (Figure 7G).

The *Pyramimonas sp.* genotype represented by **RCC5252** is formed by two IC strains from samples taken at 20 m depth at the peak of the bloom on different sampling days (Supplemntary Data S1). The representative sequence is 100% similar to that of the Arctic strain *Pyramimonas* sp. RCC1987. These strains were lost and no images are available.

*Pyramimonas australis* **RCC5269** strain from IC water has a sequence matching with 100% similarity that of *P. australis* (GenBank AJ404886) from the subgenus *Trichocystis*, an Antarctic species described based on light/electron microscopy, nuclear-encoded small-subunit ribosomal DNA and chloroplast-encoded *rbc*L gene sequences, but with no representative sequence from cultures until now (Moro et al., 2002). Cells are pear-like to almost oval, ~ 10 *µ*m long and 6 *µ*m wide with four flagella (Figure 7E).

*Pyramimonas sp.* **RCC5483** strain was recovered from IC surface waters during the pre-bloom phase and its sequence shares 100% similarity with that of the Arctic strain RCC669 (Figure 8). This strain was lost and no images are available.

*Pyramimonas sp.* **RCC5453** was isolated from an IC ice core sample during the pre-bloom phase and its sequence matches with 99.7% similarity that of the Arctic strain *Pyramimonas* sp. RCC1987. Cells are pear-like to round, from 4 to 7 *µ*m long and with four flagella (Figure 7K).

#### Cryptophyta

The *Rhodomonas* sp. genotype represented by **RCC5246** contains 14 strains collected from IC water and ice samples (Supplementary Data S1). Their representative sequence matches with 100% similarity those of the *Rhodomonas* sp. strains CCMP2045 and CCMP2293, both from Baffin Bay (Figure 6A). Cells are ~ 10 *µ*m long and 5 *µ*m wide with a prominent pyrenoid (Figure 7A). This genus is frequently observed in Arctic waters (Lovejoy et al., 2002), being abundant in the subsurface chlorophyll maximum (Joo et al., 2012) or associated with sea-ice (Niemi et al., 2011).

The *Rhodomonas* sp. genotype represented by **RCC5610** groups 6 strains, 4 of which were isolated from AM water samples (Supplementary Data S1). Its representative sequence has low similarity to that of **RCC5246** (96.8%) and is 100% similar to other Arctic strains such as RCC2020 and *Rhodomonas marina* SCCAP K-0435 from Denmark (Figure 6A), a species associated with sea-ice (Niemi et al., 2011). Cell length is ~ 18 *µ*m with a ventral to dorsal width ~ 8 *µ*m, two flagella, and a clearly visible furrow with rows of ejectisomes (Figure 7R).

#### Haptophyta

*Pseudohaptolina sorokinii* strain (**RCC5270**) was retrieved from IC water during the peak of the bloom. Its sequence shares 100% similarity with that of the recently described *P. sorokinii* (Orlova et al., 2016) strain PsAB2015 collected from coastal, under-ice water and 99.7% with the strain *P. arctica* CCMP 1204 (Figure 6C). Cells are round to oblong, ~ 17 *µ*m in length and 12 *µ*m in width. The two flagella have almost the same length as the cell, with a shorter haptonema (Figure 7F).

The *Isochrysis* sp. genotype represented by **RCC5486** contains three strains, all retrieved from IC ice core samples. Their sequence shared low similarity to any other cultured strain in the GenBank database, matching with 99.1% identity the *Isochrysis nuda* strain RCC3686 and 98.8% *Isochrysis galbana* strain 24-25B5 (Figure 6C). Cells are solitary, round to oval, ~ 6 *µ*m long and 5 *µ*m wide. The nucleus, stigma and two 7 *µ*m flagella can be observed (Figure 7L). Although mainly isolated from coastal and estuarine environments (Bendif et al., 2013), this genus has also been reported as characteristic of sea-ice environments (Majaneva et al., 2017).

#### Alveolata (Dinophyta)

The *Biecheleria cincta* genotype represented by **RCC5518** has three strains, all from AM water samples at 20 m depth during the bloom-development phase (Supplementary Data S1). Sequences from this genotype are related with 100% identity to the Arctic isolate RCC2013 *Biecheleria cincta* (Figure 6), a cosmopolitan species found also in polar waters (Balzano et al., 2012a), with reported mixotrophic behaviour (Kang et al., 2011). Cells are ~ 10 *µ*m wide with irregular shaped chloroplasts (Figure 7O).

The sequence of *Biecheleria* sp. (**RCC5522**), collected in the same sample as the *B. cincta* **RCC5518** genotype, differed by only one base pair from the sequence of RCC5518, branching with *B. brevisulcata* strain trd276-kt from freshwater (Figure 6). Cells are spherical to oval, ~ 8 *µ*m long and 6 *µ*m wide with irregularly shaped chloroplasts (Figure 7P).

### Culture diversity according to isolation source and method

#### Ice Camp

A total of 187 strains were isolated from Ice Camp samples, 110 from the water and 77 from the ice (Supplementary Data S1). Diatoms dominated isolates from all phases of the bloom (pre-bloom, boom-development and bloom-peak, see Material and Methods section), although the diversity and number of strains varied (Figure 9). During the pre-bloom phase, 28 strains were recovered from the ice and 10 from the water belonging to six classes (Figure 9). This phase was dominated by Bacillariophyceae, mainly *Nitzschia* sp. and *F. cylindrus* (Figure 9, Supplementary Data S1). Eight out of the eleven Mediophyceae strains belonged to the genus *Thalassiosira*. Strains from Pyramimonadales, Prymnesio-phyceae, Pelagophyceae and Mamiellophyceae were also retrieved during prebloom. The bloom-development phase yielded 50 strains from seven classes. New taxa not isolated during the first phase appeared, including one strain of Pedinel-lales (Dictyochophyceae) from an ice sample, and 7 strains of Cryptophyceae assigned to *Rhodomonas* sp., all from water samples (Figure 9, Supplementary Data S1). More strains were retrieved during the bloom-peak than the other two phases combined (99), and eleven classes were isolated. In contrast to the previous two phases, strains retrieved from water were more diverse than from the ice (Figure 9). Chlorophyceae were represented by *Chlamydomonas* sp., from both water and ice samples. One strain of the Prymnesiophyceae *P. sorokinii* and the Chrysophyceae strains *D. faculiferum* and *Spumella* sp. were only retrieved during this phase (Supplementary Data S1). With respect to diatoms, this phase was marked by an increase in Mediophyceae, particularly from the genera *Chaetoceros* and *Attheya* (Supplementary Data S1).

**Figure 9.**
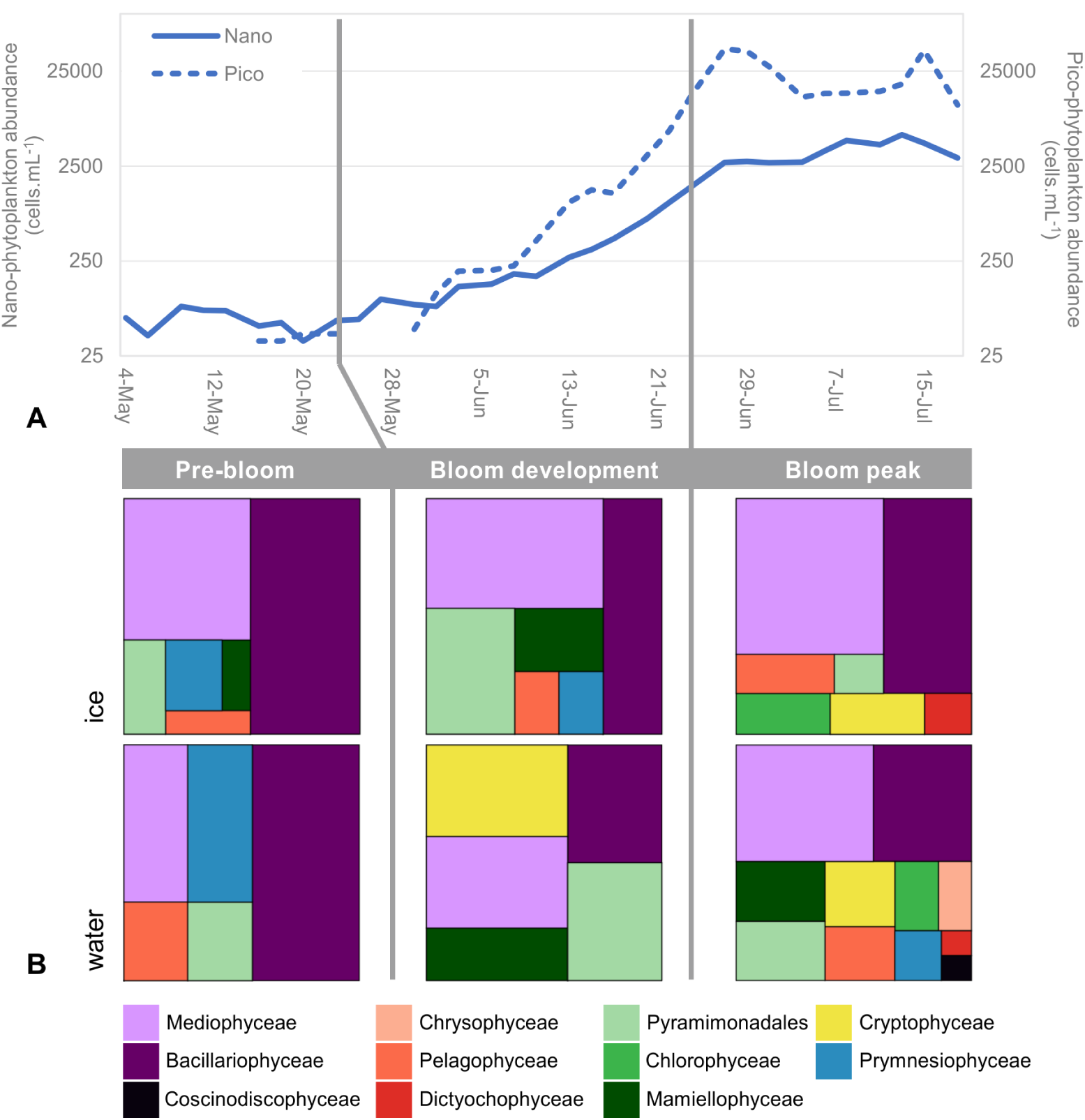
Evolution of culture diversity during the bloom. A) Abundance of pico-(dashed line, right axis) and nano-phytoplankton (solid lines, left axis) measured by flow cytometry at 10 m depth at the Ice Camp location. Phases of the bloom: pre-bloom (May 4 to 23), bloom-development (May 24 to June 22) and bloom-peak (June 23 to July 18). B) Treemaps showing the distribution of strains by class during the different phases of the bloom for the water and ice samples.

#### Amundsen cruise

89 strains were isolated from Amundsen cruise water samples, of which 81 were diatoms. Although some stations were dominated by Bacillariophyceae, such as station G102 (Figure S2), the majority of the strains belonged to the Mediophyceae, particularly *Attheya* sp. and *C. neogracilis* (Supplementary Data S1). Only two strains of the Coscinodiscophyceae genus *Actinocyclus* sp. were recovered, both from surface waters at the same station (G713). Few non-diatom strains were isolated. The only station with Cryptophyceae representatives was AM1, from which four *Rhodomonas* strains were isolated, all from the same sample and with the same isolation method (Figure S2). Four Dinophyceae strains (*Biecheleria* sp.) were retrieved from station G204. One strain of *Spumella* sp. was recovered from G110 and two *Pyramimonas* sp. strains from G512 and G707 (Figure S2, Supplementary Data S1).

#### Isolation method

Serial dilution yielded most genotypes (18), followed by FCM cell sorting (14) and single cell pipetting (7). Eighteen genotypes had representatives isolated by more than one technique (Figure S3). Among diatoms, Bacillariophyceae and Mediophyceae were retrieved by the three isolation methods, but Coscinodiscophyceae were isolated only by single cell pipetting (Figure S4A to C). Specifically, *Arcocellulus* sp. and *E. groenlandica* were retrieved only by flow cytometry sorting, while *B. bathyomphala*, *Coscinodiscus* sp. and *Actinocyclus* sp. strains came only from single cell pipetting. The strains isolated only by serial dilution included *Sellaphora* sp., *Skeletonema* sp. and *Stauroneis* sp. (Supplementary Data S1). For non-diatoms, flow cytometry cell sorting was the technique which retrieved the highest diversity at the class level (Figure S4F). *D. faculiferum*, *Mantoniella* sp., *Biecheleria* sp., and *P. sorokinii* were only obtained by this technique. *B. prasinos*, *M. baffinensis* and *Spumella* sp. were recovered only by serial dilution, as well as nine out of ten *M. polaris* strains.

## Discussion

### Novel diversity

Half of the strains in this study were retrieved using FCM cell sorting, reflecting previous reports on the efficiency of this isolation technique (Marie et al., 2017). The use of other techniques helped to increase the diversity of taxa successfully cultured, as 68% of genotypes were obtained by a single isolation method, confirming previous work in the Arctic and other marine systems (Balzano et al., 2012; Le Gall et al., 2008). For instance, although only 12% of strains originated from single cell pipetting, Coscinodiscophyceae were only retrieved by this technique, as well as three of four *Thalassiosira* genotypes. Serial dilution yielded 38% of the strains and was particularly successful for retrieving picoplanktonic Mamiellophyceae. In fact, at the early stages of isolate characterization (before screening and dereplication), 60 picoplankton strains were established by this technique, compared to only one by cell sorting and none by single cell pipetting. Among the genotypes retrieved by more than one isolation method were some well known Arctic taxa such as *A. septentrionalis*, *C. neogracilis* and *F. cylindrus*. Of the 57 retrieved genotypes, 32 could not be assigned at the species level and

6 at the genus level. Some species cannot be reliably determined by 18S rRNA sequencing alone, like *T. rotula*, *A. cornucervis* or *C. neogracilis* that may display cryptic diversity. In such cases accurate determination would usually require the use of alternative gene markers such as 28S rRNA or ITS(Balzano et al., 2017), or there may be morphological characters that distinguish the species. For example, the closely related species *A. septentrionalis* and *A. longicornis* cannot be discriminated by 18S rRNA (Rampen et al., 2009), but the latter can be distinguished morphologically by its characteristic long horns.

Of the diversity cultured in this study, pennate diatoms contained the most candidates for novel taxa (i.e. closely related only to environmental sequences). The five genotypes affiliated to *Nitzschia* sp. were not closely related to any existing sequenced strain. For example, the *Nitzschia* RCC5458 strain isolated from the ice branched apart from other *Nitzschia* genotypes with high bootstrap support (95%), with only 98% similarity to a strain from the Red Sea. Also retrieved from the ice, *Cylindrotheca* RCC5303 grouped with *C. closterium* in a moderately supported clade (72%), apparently forming a new lineage within the *C. closterium* species complex (98% similarity). Other pennate diatoms with low sequence identity matches to existing strains included Naviculales sp. RCC5387 and *Sellaphora* sp. RCC5460. With respect to centric diatoms, *Coscinodiscus* RCC5319 had the greatest dissimilarity to any existing strain sequence (95% identity), grouping with moderate bootstrap support (80%) with *C. radiatus* from the North Pacific, a species previously reported from Baffin Bay (Lovejoy et al., 2002). Unfortunately, this strain was lost before morphological analysis was undertaken. *C. decipiens* RCC5606 is interesting in that it is clearly distinguishable from the closely related *C. decipiens* RCC1997 from the Beaufort Sea (99.8% similarity) (Balzano et al., 2017) and differs from the original description (Hasle and Syvertsen, 1997) by its prominently curved chains.

Among green algae, the newly described Arctic species *M. baffinensis* (from RCC5418) and *M. beaufortii* Yau et al. (2018)), as well as the other *Mantoniella*-related strains from this work that were lost (RCC5273 and RCC5301), suggest that this genus is more diverse than other Arctic Mamiellophyceae and hosts several rare species that are not often revealed in environmental sequencing data. The Mamiellophyceae *B. prasinos* (RCC5417) strain that was isolated from ice is, to the best of our knowledge, the only available Arctic isolate of this very ubiqui-tous species (Tragin and Vaulot, 2019). This will offer interesting perspectives in terms of genome sequencing and physiological experiments, as this strain might correspond to a new cold-adapted ecotype.

The *Isochrysis* sp. strains that originated from the ice are not closely related to any polar strain or environmental sequence, potentially representing a new cold-adapted genotype. The retrieval of only one dinoflagellate species, *Biecheleria cincta* (previously *Woloszynskia* (Balzano et al., 2012) is at odds with the known diversity of dinoflagellates in the Arctic (Bachy et al., 2011; Onda et al., 2017) and especially in Baffin Bay (Lovejoy et al., 2002). Another extensive Arctic culture isolation effort yielded a similar result (Balzano et al., 2012), indicating the need for alternative isolation methods to overcome this bias.

### Change in diversity during bloom development

The strains recovered were more numerous and more diverse during the bloom itself when sea-ice melted. During the two preliminary phases of the bloom (pre-bloom and bloom-development) the highest strain diversity originated from ice samples. A shift occurred as the bloom became established and the water column samples yielded more strain diversity. There was an increase in flagellate strains isolated from water during the bloom, going from 3 during the pre-bloom period to 33 at its peak. Flagellate dominated communities have been reported in late summer in northern Baffin Bay and the Beaufort Sea (Tremblay et al., 2009). During pre-bloom, flagellates isolated from water samples belonged to only two classes (Pelagophyceae and Pyramimonadales), compared to seven classes during later phases. *Chlamydomonas* (Chlorophyceae), a genus usually associated with freshwater environments, were only isolated in July when ice melting accelerated, lowering the salinity of surface waters. All *Micromonas* and most *Pyramimonas* strains (20 out of 24) were also isolated from the two later phases of the bloom. Both genera have been documented in abundance in lower salinity, summer Arctic waters (Balzano et al., 2012; Not et al., 2005), although higher *M. polaris* abundance has been associated with both pre-bloom and post-bloom stages (Marquardt et al., 2016; Meshram et al., 2017), thriving in both nutrient-replete and nutrient-deplete conditions (Balzano et al., 2012). Flow cytometry data showed a peak in picoplankton abundance preceding that of nanoplankton (Figure 9), a pattern that has previously been observed (Sherr et al., 2003). One *M. polaris* strain was retrieved from an ice core sample during bloom-development, confirming previous studies using high throughput sequencing that have shown that *M. polaris* is part of both Arctic (Comeau et al., 2013) and subarctic sympagic communities (Belevich et al., 2018). *Pyramimonas* cell abundance in the Baffin Bay region during summer is exceptionally high compared to other Arctic domains such as the Bering, Chukchi and Beaufort Seas (Crawford et al., 2018), where it seems to be also fairly diverse (Balzano et al., 2012). Pyramimonadales were indeed the third most represented class in the present study, from both water and ice samples. Ochrophyta strains associated with heterotrophic or mixotrophic behavior such as *Spumella*, *Dinobryon* (Unrein et al., 2010) and Pedinellales (Piwosz and Pernthaler, 2010) were only isolated during the bloom-peak, which might be related to a competitive advantage under nitrogen deprivation in surface waters as the spring bloom develops.

Diatoms play a major role in sympagic assemblages (Mundy et al., 2011), and a pennate dominated community (Comeau et al., 2013) is considered a mature state of the successional stages during ice formation (Kauko et al., 2018; Niemi et al., 2011), when centric diatoms are found in lower numbers (Olsen et al., 2017). *Navicula* and *Nitzschia* representatives thrive in high abundance in the high salinity brine channels (Johnsen et al., 2018; Rózanska et al., 2009). In the present study, eight out of the sixteen genotypes retrieved solely from ice were pennate di-atoms, including two *Navicula* and two *Nitzschia* species. As the ice melts and the bloom develops in the Arctic pelagic environment, bigger cells prosper, including centric diatoms such as *T. nordenskioeldii*, *T. antarctica* var. *borealis* and/or the smaller-sized *C. gelidus* (Booth et al., 2002; Horvat et al., 2017). The relevance of the pelagic environment to centric diatoms was demonstrated by the Bacillar-iophyceae/Coscinodiscophyceae genera recovered solely from the water column, including *Skeletonema*, *Shionodiscus* and *Actinocyclus*. Although *Thalassiosira* strains were isolated from the first phase of the bloom, including ice samples, *C. gelidus* was only retrieved from mid-June onwards. *C. gelidus* has been often reported in the Arctic (Ardyna et al., 2017; Johnsen et al., 2018), in particular following *Thalassiosira* blooms (Booth et al., 2002). *C. neogracilis* strains alone comprised 12% of all strains and were retrieved from all phases of the bloom, from ice and surface waters down to 35 meters. The wide spatial and temporal range from which this species was retrieved attests for its ubiquity and importance in this environment.

## Conclusion

Ice, under-ice and open water Arctic phytoplankton communities differ in diversity, biomass, growth rate and tolerance to environmental conditions (Arrigo et al., 2012). Similarly, different types of ice provide different substrates, and therefore harbor different communities (Comeau et al., 2013; Majaneva et al., 2017). The same is true for the stages of ice formation (Kauko et al., 2018; Olsen et al., 2017). In the present work, ice core samples yielded most of the novel taxa, for all groups from diatoms to green algae. It is important that culturing efforts continue in the Arctic, as ongoing and predicted loss in ice coverage and thickness (Perovich and Richter-Menge, 2009) will certainly impact plankton diversity, dynamics and community structure (Blais et al., 2017; Comeau et al., 2011; Horvat et al., 2017). As the diversity within culture collections improves to reflect the complexity of the environment, the increased amount of validated reference sequences will help scientists to better access eukaryotic plankton distribution patterns across the Arctic. In addition, the availability of polar strains will enable experimental studies to observe physiological and metabolic impacts of current changes such as global warming on polar phytoplankton communities.

## Contributions

Contributed to conception and design: DV, ALS, IP

Contributed to acquisition of data: CGR, ALS, PG,FLG, DM, MT, IP, DV

Contributed to analysis and interpretation of data: CGR, ALS, DV

Drafted and/or revised the article: CGR, ALS, IP, DV

Approved the submitted version for publication: CGR, ALS, DV

## Acknowledgments

We thank the whole Green Edge team, as well as the Amundsen crew, for the help they provided at all stages of this project. Special thanks to Marie-Hélène Forget and Joannie Ferland for the Ice Camp logistics.

## Data accessibility statement

Strains have been deposited to the Roscoff Culture Collection (http://www.roscoff-culture-collection.org) under numbers RCC5197 to RCC5612 and sequences to Genbank under accession numbers MH764681:765044.

## Supplementary material

Supplementary data are available on GitHub at https://github.com/vaulot/Paper-2019-Ribeiro-GE-cultures

Supplementary Data S1: File GE_cultures_Tables.xlsx. Sheet Data S1. Strains collected during GE campaign, including both Amundsen an Ice Camp samples: RCC and GenBank accession number, taxonomy, respective clusters, sampling substrate, depth and date, geographic coordinates and isolation method.

Supplementary Data S2: File GE_cultures_Tables.xlsx. Sheet Data S2. Best BLAST hit for representative 18S rRNA sequences from each genotype against all GenBank sequences, PR^2^ sequences Guillou et al. (2013) and sequences from cultured strains.

**Figure S1.**
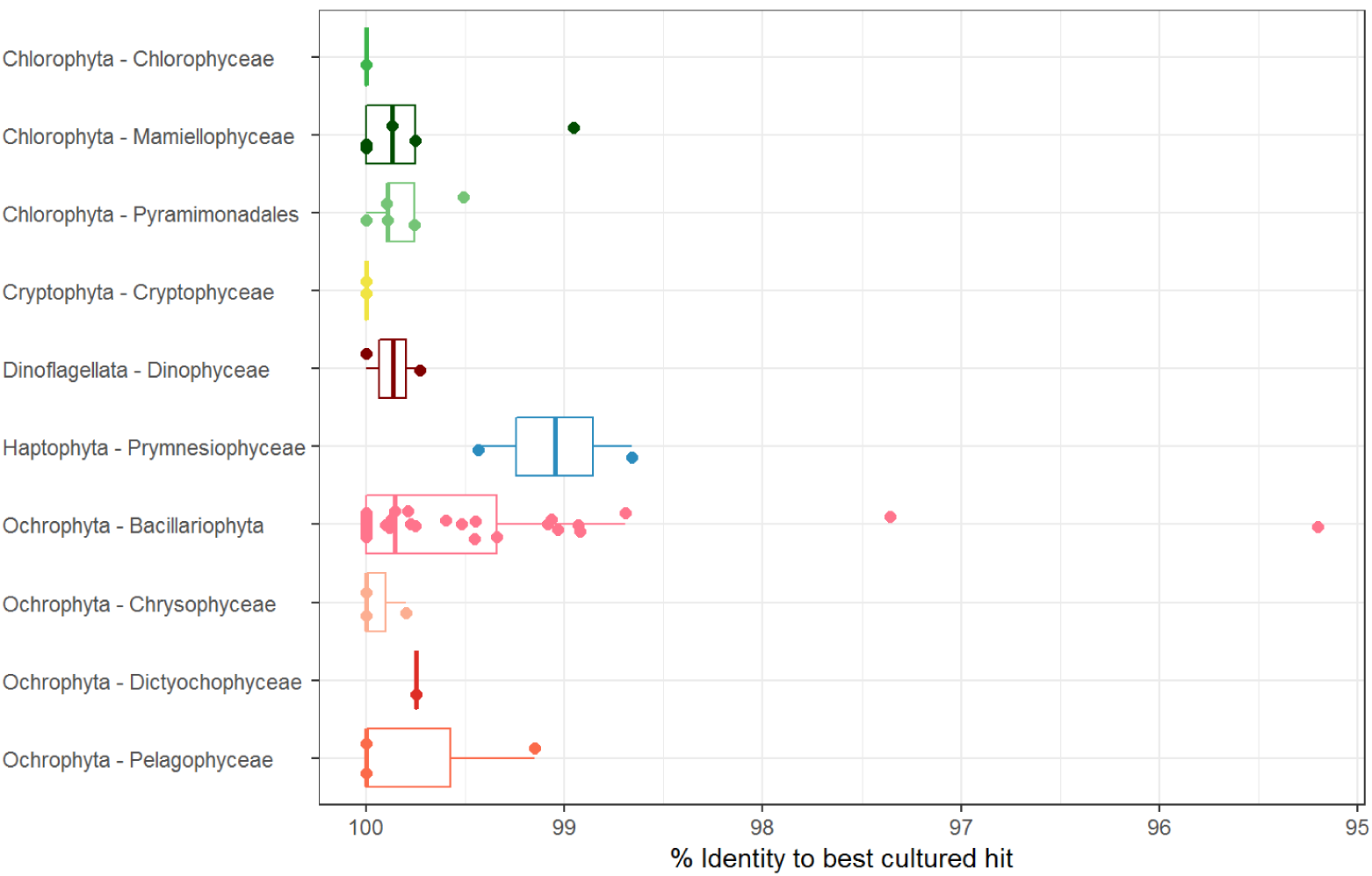
Novelty of genotypes. Percentage of similarity of genotype representative 18S rRNA sequence to best BLAST hit from GenBank (see Supplementary Data S2).

**Figure S2.**
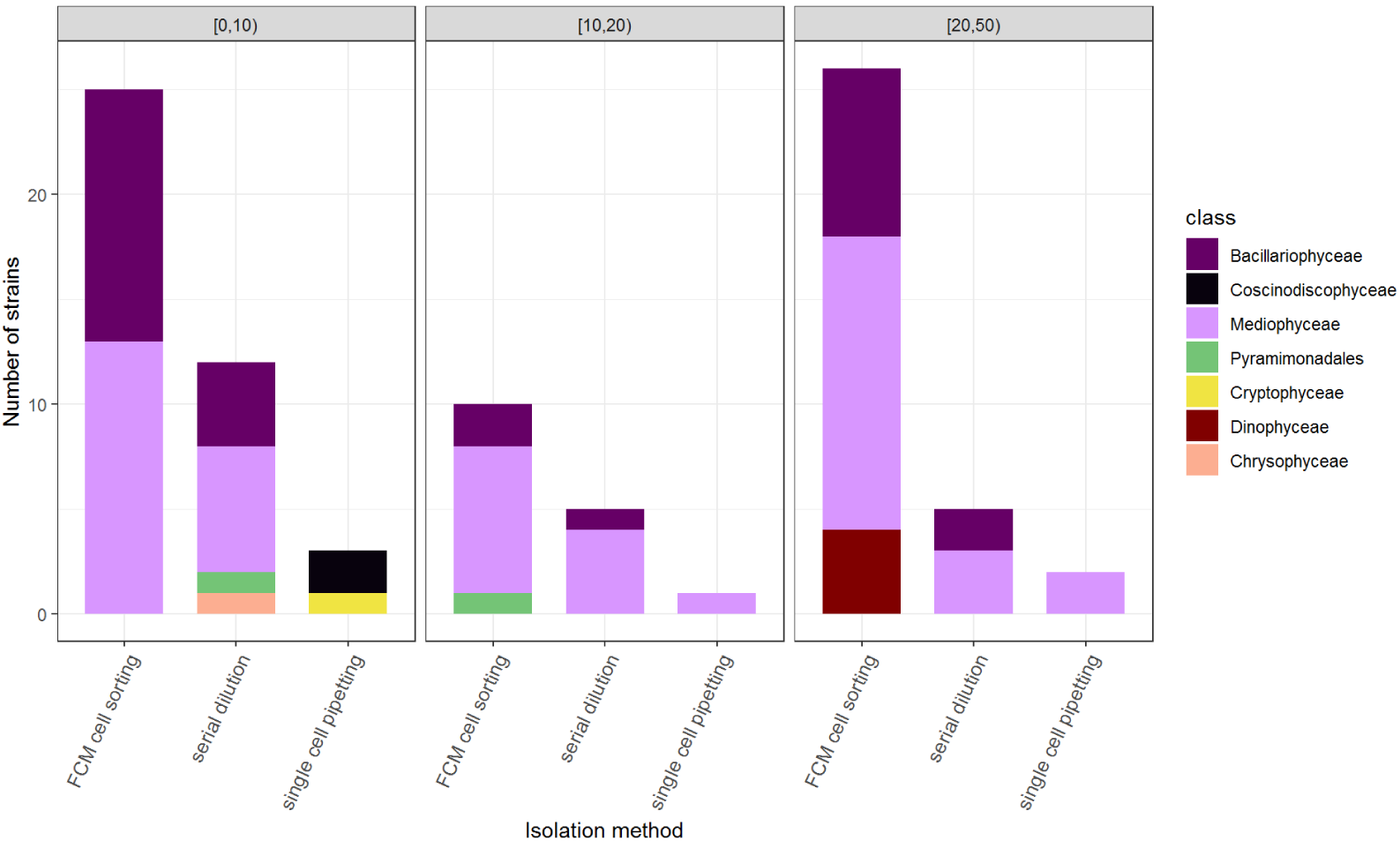
Strains from Amundsen cruise as a function of isolation method and depth. Strain class distribution for the Amundsen cruise separated according to the method of isolation (cell sorting, serial dilution and single cell isolation) and sampling depth range.

**Figure S3.**
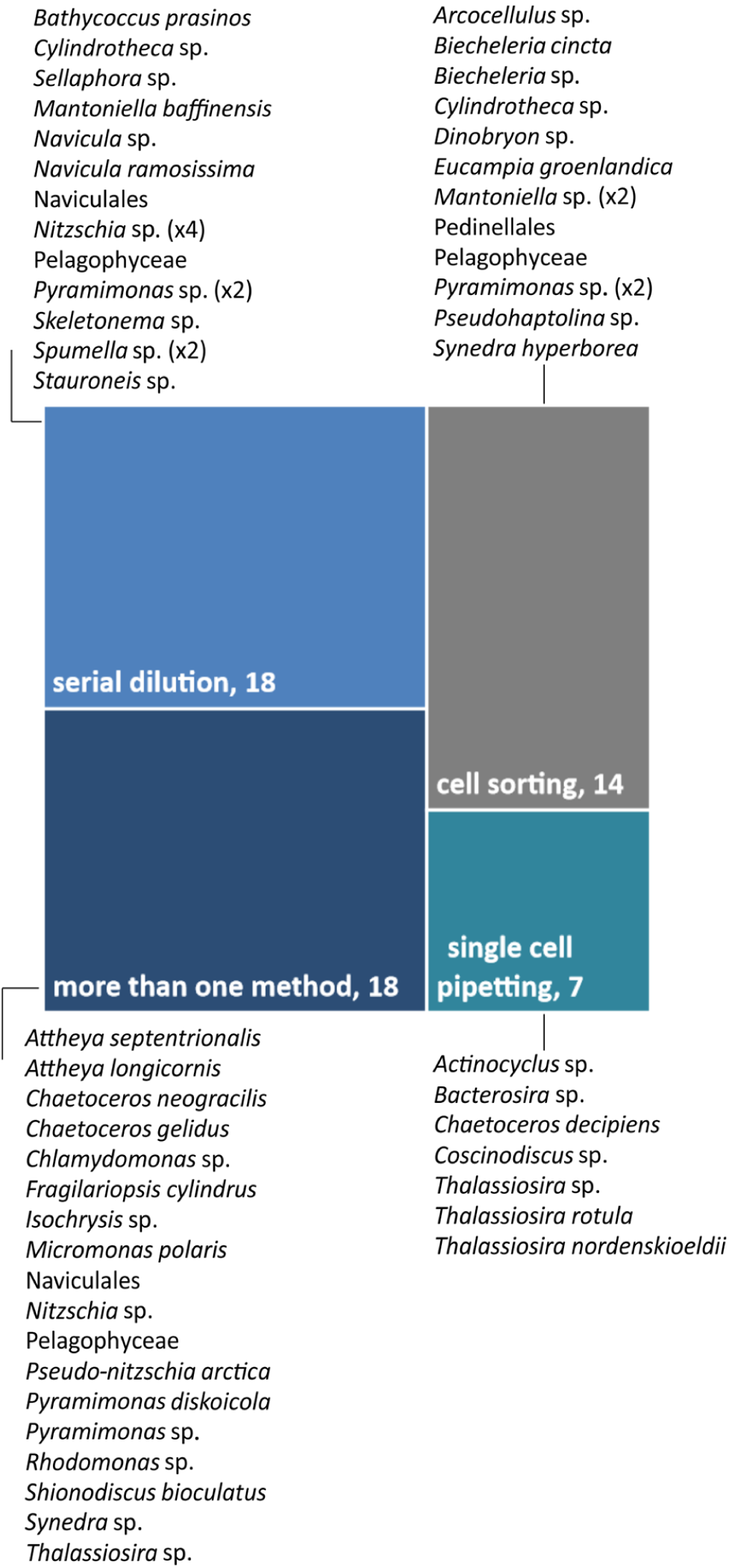
Genotype as a function of isolation method. Treemap of the number of strains isolated as function of the isolation method.

**Figure S4.**
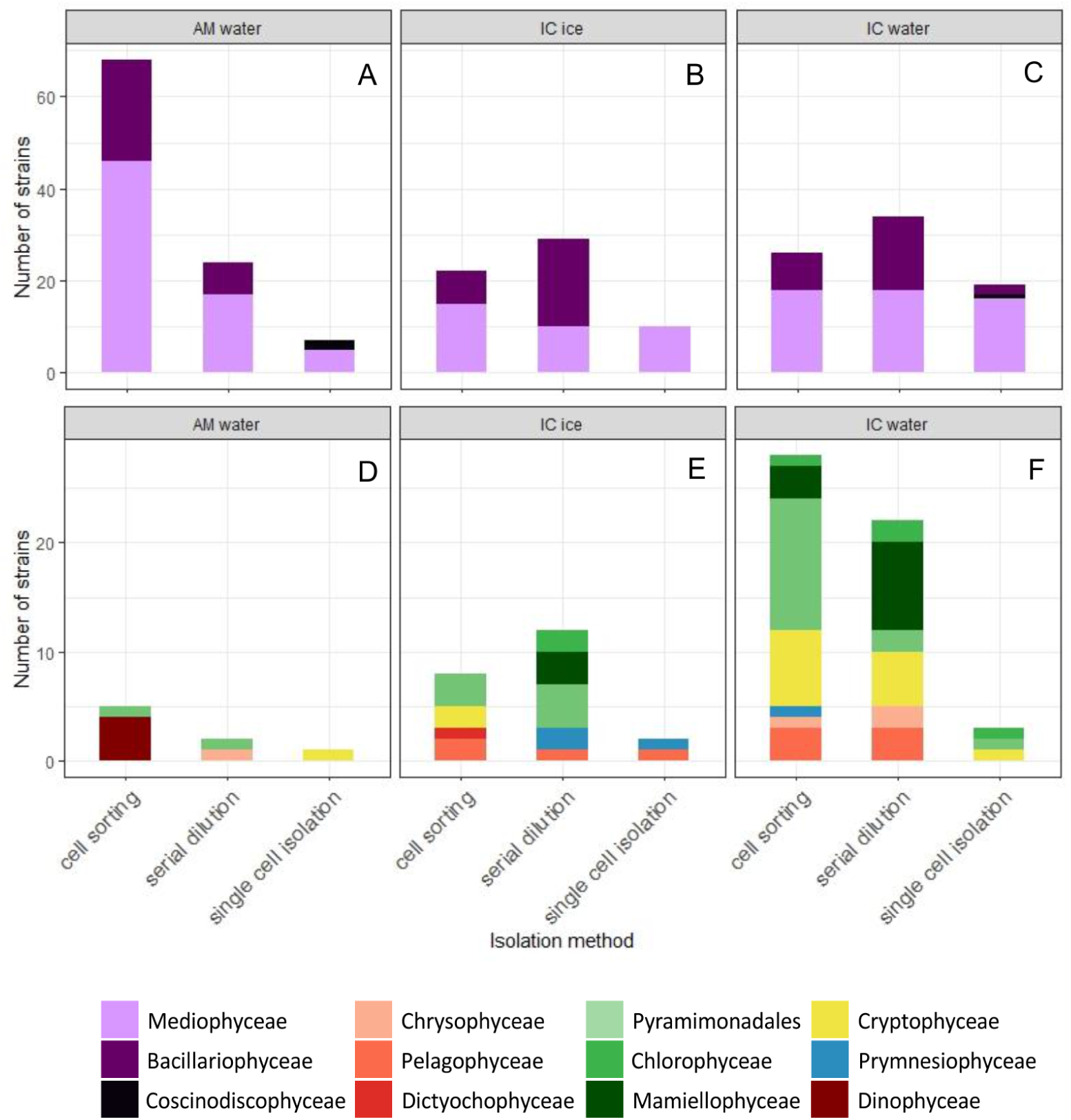
Strains as a function of isolation method and substrate. Strains class distribution separated according to the method of isolation (cell sorting, serial dilution and single cell isolation) and sampling substrate: water samples from the Amundsen cruise, and water and ice samples from the Ice Camp for diatoms (top panels) and non-diatoms (bottom panels).

## References

Ardyna M, Babin M, Devred E, Forest A, Gosselin M, et al. 2017. Shelf-basin gradients shape ecological phytoplankton niches and community composition in the coastal Arctic Ocean (Beaufort Sea). Limnology and Oceanography 62(5): 2113–2132. doi:10.1002/lno.10554.

Arrigo KR, Perovich DK, Pickart RS, Brown ZW, van Dijken GL, et al. 2012. Massive Phytoplankton Blooms Under Arctic Sea Ice. Science 336(6087): 1408–1408. doi:10.1126/science.1215065.

Arrigo KR, Perovich DK, Pickart RS, Brown ZW, van Dijken GL, et al. 2014. Phytoplankton blooms beneath the sea ice in the Chukchi sea. Deep Sea Research Part II: Topical Studies in Oceanography 105: 1–16. doi:10.1016/j.dsr2.2014.03.018.

Arrigo KR, van Dijken G, Pabi S. 2008. Impact of a shrinking Arctic ice cover on marine primary production. Geophysical Research Letters 35(19): L19603. doi:10.1029/2008GL035028.

Årthun M, Eldevik T, Smedsrud LH, Skagseth Ø, Ingvaldsen RB. 2012. Quantifying the Influence of Atlantic Heat on Barents Sea Ice Variability and Retreat. Journal of Climate 25(13): 4736–4743. doi:10.1175/JCLI-D-11-00466.1.

Assmy P, Ehn JK, Fernández-méndez M, Hop H, Katlein C, et al. 2013. Floating Ice-Algal Aggregates below Melting Arctic Sea Ice. PLoS ONE 8(10): 1–13. doi:10.1371/journal.pone.0076599.

Assmy P, Fernández-Méndez M, Duarte P, Meyer A, Randelhoff A, et al. 2017. Leads in Arctic pack ice enable early phytoplankton blooms below snow-covered sea ice. Scientific Reports 7: 40850. doi:10.1038/srep40850.

Bachy C, Lopez-Garcia P, Vereshchaka A, Moreira D. 2011. Diversity and vertical distribution of microbial eukaryotes in the snow, sea ice and seawater near the North Pole at the end of the polar night. Frontiers in Microbiology 2: 1–12. doi:10.3389/fmicb.2011.00106.

Balzano S, Gourvil P, Siano R, Chanoine M, Marie D, et al. 2012. Diversity of cultured photosynthetic flagellates in the northeast Pacific and Arctic Oceans in summer. Biogeosciences 9: 4553–4571. doi:10.5194/bg-9-4553-2012.

Balzano S, Marie D, Gourvil P, Vaulot D. 2012. Composition of the summer photosynthetic pico and nanoplankton communities in the Beaufort Sea assessed by T-RFLP and sequences of the 18S rRNA gene from flow cytometry sorted samples. The ISME journal 6: 1480–1498. doi:10.1038/ismej.2011.213.

Balzano S, Percopo I, Siano R, Gourvil P, Chanoine M, et al. 2017. Morphological and genetic diversity of Beaufort Sea diatoms with high contributions from the *Chaetoceros neogracilis* species complex. Journal of Phycology 53: 161– 187. doi:10.1111/jpy.12489.

Belevich TA, Ilyash LV, Milyutina IA, Logacheva MD, Goryunov DV, et al. 2018. Photosynthetic Picoeukaryotes in the Land-Fast Ice of the White Sea, Russia. Microbial Ecology 75: 582–597. doi:10.1007/s00248-017-1076-x.

Bendif EM, Probert I, Schroeder DC, de Vargas C. 2013. On the description of *Tisochrysis lutea* gen. nov. sp. nov. and Isochrysis nuda sp. nov. in the Isochrysidales, and the transfer of *Dicrateria* to the Prymnesiales (Haptophyta). Journal of Applied Phycology 25: 1763–1776. doi:10.1007/s10811-013-0037-0.

Blais M, Ardyna M, Gosselin M, Dumont D, Simon B, et al. 2017. Contrasting interannual changes in phytoplankton productivity and community structure in the coastal Canadian Arctic Ocean. Limnology and Oceanography 62: 2480– 2497. doi:10.1002/lno.10581.

Boenigk J, Pfandl K, Stadler P, Chatzinotas A. 2005. High diversity of the ‘*Spumella*-like’ flagellates: an investigation based on the SSU rRNA gene sequences of isolates from habitats located in six different geographic regions. Environmental Microbiology 7(5): 685–697. doi:10.1111/j.1462-2920.2005.00743.x.

Booth BC, Larouche P, Bélanger S, Klein B, Amiel D, et al. 2002. Dynamics of *Chaetoceros socialis* blooms in the North Water. Deep-Sea Research Part II: Topical Studies in Oceanography 49(22-23): 5003–5025. doi:10.1016/S0967-0645(02)00175-3.

Brown ZW, Arrigo KR. 2013. Sea ice impacts on spring bloom dynamics and net primary production in the Eastern Bering Sea. Journal of Geophysical Research: Oceans 118(1): 43–62. doi:10.1029/2012JC008034.

Campbell K, Mundy CJ, Belzile C, Delaforge A, Rysgaard S. 2017. Seasonal dynamics of algal and bacterial communities in Arctic sea ice under variable snow cover. Polar Biology 41(1): 41–58. doi:10.1007/s00300-017-2168-2.

Chamnansinp A, Li Y, Lundholm N, Moestrup Ø. 2013. Global diversity of two widespread, colony-forming diatoms of the marine plankton, Chaetoceros socialis (syn. C. radians) and Chaetoceros gelidus sp. nov. Journal of phy 49: 1128–1141. doi:10.1111/jpy.12121.

Cleve PT. 1896. Diatoms from Baffin Bay and Davis Strait. Bihang till Kongliga Svenska Vetenskaps-Akademiens Handlingar 22: 1–22.

Comeau AM, Li WKW, Tremblay JE, Carmack EC, Lovejoy C. 2011. Arctic Ocean Microbial Community Structure before and after the 2007 Record Sea Ice Minimum. PLoS ONE 6(11): 1–12. doi:10.1371/journal.pone.0027492.

Comeau AM, Philippe B, Thaler M, Gosselin M, Poulin M, et al. 2013. Protists in Arctic Drift and Land-Fast Sea Ice. Journal of Phycology 49(2): 229–240. doi:10.1111/jpy.12026.

Crawford DW, Cefarelli AO, Wrohan IA, Wyatt SN, Varela DE. 2018. Spatial patterns in abundance, taxonomic composition and carbon biomass of nano- and microphytoplankton in Subarctic and Arctic Seas. Progress in Oceanography 162(October 2016): 132–159. doi:10.1016/j.pocean.2018.01.006.

Daugbjerg N, Moestrup Ø. 1993. Four new species of *Pyramimonas* (Prasinophyceae) from arctic Canada including a light and electron microscopic description of *Pyramimonas quadrifolia* sp. nov. European Journal of Phycology 28: 3–16. doi:10.1080/09670269300650021.

Delmont TO, Murat Eren A, Vineis JH, Post AF. 2015. Genome reconstructions indicate the partitioning of ecological functions inside a phytoplankton bloom in the Amundsen Sea, Antarctica. Frontiers in Microbiology 6: 1–19. doi:10.3389/fmicb.2015.01090.

Demory D, Baudoux AC, Monier A, Simon N, Six C, et al. 2018. Picoeukaryotes of the *Micromonas* genus: sentinels of a warming ocean. The ISME Journal doi:10.1038/s41396-018-0248-0.

Duerksen SW, Thiemann GW, Budge SM. 2014. Large, Omega-3 Rich, Pelagic Diatoms under Arctic Sea Ice: Sources and Implications for Food Webs. PloS one 9(12): 1–18. doi:10.1371/journal.pone.0114070.

Eilertsen HC, Degerlund M. 2010. Phytoplankton and light during the northern high-latitude winter. Journal of Plankton Research 32(6): 899–912. doi:10.1093/plankt/fbq017.

Felsenstein J. 1985. Confidence Limits on Phylogenies: An Approach Using the Bootstrap. Evolution 39(4): 783–791. doi:10.3389/fimmu.2015.00048.

Fernández-méndez M, Olsen LM, Kauko HM, Meyer A, Rösel A, et al. 2018. Algal Hot Spots in a Changing Arctic Ocean: Sea-Ice Ridges and the Snow-Ice Interface. Frontiers in Marine Science 5: 1–22. doi:10.3389/fmars.2018.00075.

Grossmann L, Bock C, Schweikert M, Boenigk J. 2015. Small but Manifold – Hidden Diversity in “*Spumella*-like Flagellates”. Journal of Eukaryotic Microbiology 0: 1–21. doi:10.1111/jeu.12287.

Guillard RRL, Hargraves PE. 1993. *Stichochrysis immobilis* is a diatom, not a chrysophyte. Phycologia 32(3): 234–236. doi:10.2216/i0031-8884-32-3-234.1.

Guillou L, Bachar D, Audic S, Bass D, Berney C, et al. 2013. The Protist Ribosomal Reference database (PR^2^): a catalog of unicellular eukaryote Small Sub-Unit rRNA sequences with curated taxonomy. Nucleic Acids Research 41(D1): D597–D604. doi:10.1093/nar/gks1160.

Guindon S, Dufayard JF, Lefort V, Anisimova M, Hordijk W, et al. 2010. New algorithms and methods to estimate maximum-likelihood phylogenies: assessing the performance of PhyML 3.0. Systematic Biology 59(3): 307–321. doi:10.1093/sysbio/syq010.

Harðardóttir S, Lundholm N, Moestrup Ø, Nielsen TG. 2014. Description of *Pyramimonas diskoicola* sp. nov. and the importance of the flagellate Pyramimonas (Prasinophyceae) in Greenland sea ice during the winter–spring transition. Polar Biology 37(10): 1479–1494. doi:10.1007/s00300-014-1538-2.

Hasle GR, Heimdal BR. 1968. Morphology and distribution of the marine centric diatom *Thalassiosira antarctica* Comber. Journal of the Royal Microscopical Society 88(3): 357–369.

Hasle GR, Medlin LK, Syvertsen EE. 1994. *Synedropsis* gen. nov., a genus of araphid diatoms associated with sea ice. Phycologia 33(4): 248–270. doi:10.2216/i0031-8884-33-4-248.1.

Hasle GR, Syvertsen EE. 1997. Marine diatoms, in Tomas CR, ed., Identifying Marine Phytoplankton. San Diego, California: Academic Press: pp. 5–385. ISBN 9780126930184.

Horvat C, Jones DR, Iams S, Schroeder D, Flocco D, et al. 2017. The frequency and extent of sub-ice phytoplankton blooms in the Arctic Ocean. Science Advances 3: 1–8. doi:10.1126/sciadv.1601191.

Johnsen G, Norli M, Moline M, Robbins I, von Quillfeldt C, et al. 2018. The advective origin of an under-ice spring bloom in the Arctic Ocean using multiple observational platforms. Polar Biology 41(6): 1197–1216. doi:10.1007/s00300-018-2278-5.

Joli N, Monier A, Logares R, Lovejoy C. 2017. Seasonal patterns in Arctic prasinophytes and inferred ecology of *Bathycoccus* unveiled in an Arctic winter metagenome. ISME Journal pp. 1–14. doi:10.1038/ismej.2017.7.

Joo HM, Lee SH, Jung SW, Dahms Hu, Lee JH. 2012. Latitudinal variation of phytoplankton communities in the western Arctic Ocean. Deep-Sea Research Part II 81–84: 3–17. doi:10.1016/j.dsr2.2011.06.004.

Kang NS, Jeong HJ, Yoo YD, Yoon EY, Lee KH, et al. 2011. Mixotrophy in the newly described phototrophic dinoflagellate *Woloszynskia cincta* from western Korean waters: Feeding mechanism, prey species and Effect of prey concentration. Journal of Eukaryotic Microbiology 58(2): 152–170. doi:10.1111/j.1550-7408.2011.00531.x.

Katsuki K, Takahashi K, Onodera J, Jordan RW, Suto I. 2009. Living diatoms in the vicinity of the North Pole, summer 2004. Micropaleontology 55(2-3): 137–170.

Kauko HM, Olsen LM, Duarte P, Peeken I, Granskog MA, et al. 2018. Algal Colonization of Young Arctic Sea Ice in Spring. Frontiers in Marine Science 5: 1–20. doi:10.3389/fmars.2018.00199.

Kearse M, Moir R, Wilson A, Stones-Havas S, Cheung M, et al. 2012. Geneious Basic: An integrated and extendable desktop software platform for the organization and analysis of sequence data. Bioinformatics 28(12): 1647–1649. doi:10.1093/bioinformatics/bts199.

Keller MD, Selvin RC, Claus W, Guillard RRL. 1987. Media for the culture of oceanic ultraphytoplankton. Journal of Phycology 23(4): 633–638. doi:10.1111/j.1529-8817.1987.tb04217.x.

Kilias ES, Nöthig EM, Wolf C, Metfies K. 2014. Picoeukaryote plankton composition off West Spitsbergen at the entrance to the Arctic Ocean. The Journal of eukaryotic microbiology 0: 1–11. doi:10.1111/jeu.12134.

Kohlbach D, Graeve M, A Lange B, David C, Peeken I, et al. 2016. The importance of ice algae-produced carbon in the central Arctic Ocean ecosystem: Food web relationships revealed by lipid and stable isotope analyses. Limnology and Oceanography 61: 2027–2044. doi:10.1002/lno.10351.

Le Gall F, Rigaut-Jalabert F, Marie D, Garczarek L, Viprey M, et al. 2008. Picoplankton diversity in the South-East Pacific Ocean from cultures. Biogeosciences 5: 203–214. doi:10.5194/bg-5-203-2008.

Lee JM, Lee JH. 2012. Morphological study of the genus *Eucampia* (Bacillariophyceae) in Korean coastal waters. Algae 27(4): 235–247. doi:10.4490/algae.2012.27.4.235.

Leeuwe MAV, Tedesco L, Arrigo KR, Assmy P, Meiners KM, et al. 2018. Microalgal community structure and primary production in Arctic and Antarctic sea ice: A synthesis. Elementa 6(4): 1–25.

Lepère C, Demura M, Kawachi M, Romac S, Probert I, et al. 2011. Whole-genome amplification (WGA) of marine photosynthetic eukaryote populations. FEMS Microbiology Ecology 76: 513–523. doi:10.1111/j.1574-6941.2011.01072.x.

Leu E, Mundy CJ, Assmy P, Campbell K, Gabrielsen TM, et al. 2015. Arctic spring awakening - Steering principles behind the phenology of vernal ice algal blooms. Progress in Oceanography 139: 151–170. doi:10.1016/j.pocean.2015.07.012.

Leu E, Søreide JE, Hessen DO, Falk-Petersen S, Berge J. 2011. Consequences of changing sea-ice cover for primary and secondary producers in the European Arctic shelf seas: Timing, quantity, and quality. Progress in Oceanography 90: 18–32. doi:10.1016/j.pocean.2011.02.004.

Li H, Yang G, Sun Y, Wu S, Zhang X. 2007. *Cylindrotheca closterium* is a species complex as was evidenced by the variations of rbcL gene and SSU rDNA. Journal of Ocean University of China 6(2): 167–174. doi:10.1007/s11802-007-0167-6.

Li WK, McLaughlin FA, Lovejoy C, Carmack EC. 2009. Smallest algae thrive as the arctic ocean freshens. Science 326: 539. doi:10.1126/science.1179798.

Lovejoy C, Legendre L, Martineau MJ, Bâcle J, Von Quillfeldt CH. 2002. Distribution of phytoplankton and other protists in the North Water. Deep-Sea Research Part II: Topical Studies in Oceanography 49: 5027–5047. doi:10.1016/S0967-0645(02)00176-5.

Lovejoy C, Massana R, Pedro C. 2006. Diversity and Distribution of Marine Microbial Eukaryotes in the Arctic Ocean and Adjacent Seas. Applied and Environmental Microbiology 72(5): 3085–3095. doi:10.1128/AEM.72.5.3085.

Lovejoy C, Vincent WF, Bonilla S, Roy S, Martineau MJ, et al. 2007. Distribution, phylogeny, and growth of cold-adapted picoprasinophytes in arctic seas. Journal of Phycology 43(1): 78–89. doi:10.1111/j.1529-8817.2006.00310.x.

Luddington IA, Lovejoy C, Kaczmarska I, Moisander P. 2016. Species-rich meta-communities of the diatom order Thalassiosirales in the Arctic and northern Atlantic Ocean. Journal of Plankton Research 38(4): 781–797. doi:10.1093/plankt/fbw030.

Lutz S, Anesio AM, Raiswell R, Edwards A, Newton RJ, et al. 2016. The biogeography of red snow microbiomes and their role in melting arctic glaciers. Nature Communications 7: 1–9. doi:10.1038/ncomms11968.

Majaneva M, Blomster J, Müller S, Autio R, Majaneva S, et al. 2017. Sea-ice eukaryotes of the Gulf of Finland, Baltic Sea, and evidence for herbivory on weakly shade-adapted ice algae. European Journal of Protistology 57: 1–15. doi:10.1016/j.ejop.2016.10.005.

Malviya S, Scalco E, Audic S, Vincent F, Veluchamy A, et al. 2016. Insights into global diatom distribution and diversity in the world’s ocean. Proceedings of the National Academy of Sciences 348(6237): 1–10. doi:10.1073/pnas.1509523113.

Marie D, Le Gall F, Edern R, Gourvil P, Vaulot D. 2017. Improvement of phytoplankton culture isolation using single cell sorting by flow cytometry. Journal of Phycology 53(2): 271–282.

Marquardt M, Vader A, Stübner EI, Reigstad M, Gabrielsen TM. 2016. Strong Seasonality of Marine Microbial Eukaryotes in a High-Arctic Fjord (Isfjorden, in West Spitsbergen, Norway). Applied and Environmental Microbiology 82(6): 1868–1880. doi:10.1128/AEM.03208-15.Editor.

Meshram AR, Vader A, Kristiansen S, Gabrielsen TM. 2017. Microbial Eukaryotes in an Arctic Under-Ice Spring Bloom North of. Frontiers in Microbiology 8: 1–12. doi:10.3389/fmicb.2017.01099.

Mock T, Robert P, Strauss J, Mcmullan M, Paajanen P, et al. 2017. Evolutionary genomics of the cold-adapted diatom *Fragilariopsis cylindrus*. Nature 541: 536–540. doi:10.1038/nature20803.

Moro I, Rocca NL, Valle LD, Moschin E, Negrisolo E, et al. 2002. *Pyramimonas australis* sp. nov. (Prasinophyceae, Chlorophyta) from Antarctica: fine structure and molecular phylogeny. European Journal of Phycology 37: 103–114.

Mundy CJ, Gosselin M, Ehn JK, Belzile C, Poulin M, et al. 2011. Characteristics of two distinct high-light acclimated algal communities during advanced stages of sea ice melt. Polar Biology 34(12): 1869–1886. doi:10.1007/s00300-011-0998-x.

Neukermans G, Oziel L, Babin M. 2018. Increased intrusion of warming Atlantic waters leads to rapid expansion of temperate phytoplankton in the Arctic. Global Change Biology pp. 1–9. doi:10.1111/gcb.14075.

Niemi A, Michel C, Hille K, Poulin M. 2011. Protist assemblages in winter sea ice: setting the stage for the spring ice algal bloom. Polar Biology 34: 1803– 1817. doi:10.1007/s00300-011-1059-1.

Not F, Massana R, Latasa M, Marie D, Colson C, et al. 2005. Late summer community composition and abundance of photosynthetic picoeukaryotes in Norwegian and Barents Seas. Limnology and Oceanography 50(5): 1677–1686.

Olsen LM, Laney SR, Duarte P, Kauko HM, Fernández-méndez M, et al. 2017. The multiyear ice seed repository hypothesis. Journal of Geophysical Research: Biogeosciences 122: 1529–1548. doi:10.1002/2016JG003668.

Onda DFL, Medrinal E, Comeau AM, Thaler M, Babin M, et al. 2017. Seasonal and Interannual Changes in Ciliate and Dinoflagellate Species Assemblages in the Arctic Ocean (Amundsen Gulf, Beaufort Sea, Canada). Frontiers in Marine Science 4(January). doi:10.3389/fmars.2017.00016.

Orlova TY, Efimova KV, Stonik IV. 2016. Morphology and molecular phylogeny of *Pseudohaptolina sorokinii* sp. nov. (Prymnesiales, Haptophyta) from the Sea of Japan, Russia. Phycologia 55(5): 506–514. doi:10.2216/15-107.1.

Percopo I, Ruggiero M, Balzano S, Gourvil P, Lundhölm N, et al. 2016. *P. arctica* sp. nov., a new cold-water cryptic Pseudo-nitzschia species within the P. pseudodelicatissima complex. Journal of Phycology 52: 184–199.

Perovich DK, Richter-Menge JA. 2009. Loss of Sea Ice in the Arctic. Annual Review of Marine Science 1(1): 417–441. doi:10.1146/annurev.marine.010908.163805.

Perrette M, Yool A, Quartly GD, Popova EE. 2011. Near-ubiquity of ice-edge blooms in the Arctic. Biogeosciences 8: 515–524. doi:10.5194/bg-8-515-2011.

Piwosz K, Pernthaler J. 2010. Seasonal population dynamics and trophic role of planktonic nanoflagellates in coastal surface waters of the Southern Baltic Sea. Environmental Microbiology 12(2): 364–377. doi:10.1111/j.1462-2920.2009.02074.x.

Pocock T, Lachance MA, Proschold T, Priscu JC, Kim SS, et al. 2004. Identification of a psychrophilic green alga from Lake Bonney Antarctica: *Chlamydomonas raudensis* ettl. (UWO 241) Chlorophyceae. Journal of Phycology 40: 1138–1148. doi:10.1111/j.1529-8817.2004.04060.x.

Poulíčková A, Špačková J, Kelly MG, Duchoslav M, Mann DG. 2008. Ecological variation within *Sellaphora* species complexes (Bacillariophyceae): Specialists or generalists? Hydrobiologia 614(1): 373–386. doi:10.1007/s10750-008-9521-y.

Poulin M, Daugbjerg N, Gradinger R, Ilyash L, Ratkova T, et al. 2011. The pan-Arctic biodiversity of marine pelagic and sea-ice unicellular eukaryotes: a first-attempt assessment. Marine Biodiversity 41(1): 13–28. doi:10.1007/s12526-010-0058-8.

Quillfeldt CHV. 2004. The diatom *Fragilariopsis cylindrus* and its potential as an indicator species for cold water rather than for sea ice. Vie et Milieu 54(2-3): 137–143.

Rampen SW, Schouten S, Elda Panoto F, Brink M, Andersen RA, et al. 2009. Phylogenetic Position of *Attheya longicornis* and *Attheya septentrionalis* (Bacillariophyta). Journal of Phycology 45(2): 444–453. doi:10.1111/j.1529-8817.2009.00657.x.

Rippka R, Coursin T, Hess W, Lichtle C, Scanlan DJ, et al. 2000. Prochlorococcus marinus Chisholm et al. 1992 subsp. pastoris subsp. nov. strain PCC 9511, the first axenic chlorophyll a2/b2-containing cyanobacterium (Oxyphotobacteria). International Journal of Systematic and Evolutionary Microbiology 50: 1833–1847. doi:10.1099/00207713-50-5-1833.

Rózanska M, Gosselin M, Poulin M, Wiktor JM, Michel C. 2009. Influence of environmental factors on the development of bottom ice protist communities during the winter – spring transition. Marine Ecology Progress Series 386: 43–59. doi:10.3354/meps08092.

Schmidt K, Brown T, Belt S, Ireland L, Taylor KW, et al. 2018. Do pelagic grazers benefit from sea ice? Insights from the Antarctic sea ice proxy IPSO25. Biogeosciences 15: 1987–2006. doi:10.5194/bg-15-1987-2018.

Sherr EB, Sherr BF, Wheeler PA, Thompson K. 2003. Temporal and spatial variation in stocks of autotrophic and heterotrophic microbes in the upper water column of the central Arctic Ocean. Deep-Sea Research Part I: Oceanographic Research Papers 50(5): 557–571. doi:10.1016/S0967-0637(03)00031-1.

Simmons MP, Bachy C, Sudek S, Van Baren MJ, Sudek L, et al. 2015. Intron invasions trace algal speciation and reveal nearly identical arctic and antarctic micromonas populations. Molecular Biology and Evolution 32(9): 2219–2235. doi:10.1093/molbev/msv122.

Simon N, Foulon E, Grulois D, Six C, Desdevises Y, et al. 2017. Revision of the Genus *Micromonas* Manton et Parke (Chlorophyta, Mamiellophyceae), of the Type Species *M. pusilla* (Butcher) Manton & Parke and of the Species *M. commoda* van Baren, Bachy and Worden and Description of two New Species Based on the Genetic and Phenotypic Characterization of Cultured Isolates. Protist 168(5): 612–635. doi:10.1016/j.protis.2017.09.002.

Terrado R, Scarcella K, Thaler M, Vincent WF, Lovejoy C. 2013. Small phytoplankton in Arctic seas: vulnerability to climate change. Biodiversity 14(1): 2–18.

Tragin M, Vaulot D. 2019. Novel diversity within marine Mamiellophyceae (Chlorophyta) unveiled by metabarcoding. Scientific Reports 9(1): 5190. doi:10.1038/s41598-019-41680-6.

Tremblay G, Belzile C, Gosselin M, Poulin M, Roy S, et al. 2009. Late summer phytoplankton distribution along a 3500 km transect in Canadian Arctic waters: Strong numerical dominance by picoeukaryotes. Aquatic Microbial Ecology 54(1): 55–70. doi:10.3354/ame01257.

Unrein F, Gasol JM, Massana R. 2010. *Dinobryon faculiferum* (Chrysophyta) in coastal Mediterranean seawater: Presence and grazing impact on bacteria. Journal of Plankton Research 32(4): 559–564. doi:10.1093/plankt/fbp150.

Vincent WF. 2010. Microbial ecosystem responses to rapid climate change in the Arctic. ISME Journal 4(9): 1089–1091. doi:10.1038/ismej.2010.108.

Whittaker KA, Rignanese DR, Olson RJ, Rynearson TA. 2012. Molecular subdivision of the marine diatom *Thalassiosira rotula* in relation to geographic distribution, genome size, and physiology. BMC Evolutionary Biology 12(209): 2–14. doi:10.1186/1471-2148-12-209.

Worden AZ, Lee JH, Mock T, Rouzé P, Simmons MP, et al. 2009. Green evolution and dynamic adaptations revealed by genomes of the marine picoeukaryotes Micromonas. Science 324: 268–272.

Yau S, Lopes dos Santos A, Eikrem W, Gérikas Ribero C, Gourvil P, et al. 2018. *Mantoniella beaufortii* and *Mantoniella baffinensis* sp. now. (Mamiellales, Mamiellophyceae), two new green algal species from the high Arctic. bioRxiv doi:10.1101/506915.

Zhu F, Massana R, Not F, Marie D, Vaulot D. 2005. Mapping of picoeucaryotes in marine ecosystems with quantitative PCR of the 18S rRNA gene. FEMS Microbiology Ecology 52: 79–92. doi:10.1016/j.femsec.2004.10.006.

